# Pan-genomic analysis of transcriptional modules across *Salmonella* Typhimurium reveals the regulatory landscape of different strains

**DOI:** 10.1101/2022.01.11.475931

**Authors:** Yuan Yuan, Yara Seif, Kevin Rychel, Reo Yoo, Siddharth Chauhan, Saugat Poudel, Tahani Al-bulushi, Bernhard O. Palsson, Anand Sastry

## Abstract

*Salmonella enterica* Typhimurium is a serious pathogen that is involved in human nontyphoidal infections. Tackling Typhimurium infections is difficult due to the species’ dynamic adaptation to its environment, which is dictated by a complex transcriptional regulatory network (TRN). While traditional biomolecular methods provide characterizations of specific regulators, it is laborious to construct the global TRN structure from this bottom-up approach. Here, we used a machine learning technique to understand the transcriptional signatures of *S. enterica* Typhimurium from the top down, as a whole and in individual strains. Furthermore, we conducted cross-strain comparison of 6 strains in serovar Typhimurium to investigate similarities and differences in their TRNs with pan-genomic analysis. By decomposing all the publicly available RNA-Seq data of Typhimurium with independent component analysis (ICA), we obtained over 400 independently modulated sets of genes, called iModulons. Through analysis of these iModulons, we 1) discover three transport iModulons linked to antibiotic resistance, 2) describe concerted responses to cationic antimicrobial peptides (CAMPs), 3) uncover evidence towards new regulons, and 4) identify two iModulons linked to bile responses in strain ST4/74. We extend this analysis across the pan-genome to show that strain-specific iModulons 5) reveal different genetic signatures in pathogenicity islands that explain phenotypes and 6) capture the activity of different phages in the studied strains. Using all high-quality publicly-available RNA-Seq data to date, we present a comprehensive, data-driven Typhimurium TRN. It is conceivable that with more high-quality datasets from more strains, the approach used in this study will continue to guide our investigation in understanding the pan-transcriptome of Typhimurium. Interactive dashboards for all gene modules in this project are available at https://imodulondb.org/ under the “*Salmonella* Typhimurium” page to enable browsing for interested researchers.

## Introduction

*Salmonella enterica* is one of the leading causes of foodborne illnesses globally^1^. Nontyphoidal *Salmonella* (NTS) is highly diverse and contains more than 2,600 serovars^2^. Among them, *S. enterica* Typhimurium is of particular interest because it has broad host specificity and poses serious challenges to public health, especially with the rise in its antibiotic resistance and the advent of new strains causing serious to life-threatening infections in sub-Saharan Africa^3^. Infections caused by Typhimurium are difficult to combat for two primary reasons. First, Typhimurium contains a wide variety of strains with different genetic and phenotypic signatures. These strains vary in virulence, persistence and response to diverse conditions, thus requiring different treatment methods. Second, Typhimurium strains use a set of intricate mechanisms to adapt to their host, develop drug resistance and enhance virulence. These mechanisms are activated by the transcriptional regulatory networks (TRNs) that coordinate gene expression under a variety of different conditions, including antibiotic treatment, starvation and stress. Well-characterized TRNs of the serovar as a whole and in individual strains would enable us to develop a better understanding of Typhimurium’s dynamic adaptation to environmental perturbations and make better predictions of clinical treatment outcomes. Gaining a deeper understanding of the TRN of *S. enterica* Typhimurium therefore holds great importance for public health.

However, efforts to characterize Typhimurium’s TRNs are mainly confined to understanding the TRNs of individual strains. Prior studies comparing transcriptomic patterns across Typhimurium strains typically focused on observing differentially expressed genes of just two strains^4^. Systems-level investigations of TRNs across more strains and over the entire serovar were hindered by inadequate analytical methods.

The increasing number of sequenced genomes has empowered pan-genomic analysis to study the genetic diversity and composition of a species. A pan-genome contains all the genes in that species, and a core genome is defined as the genes shared between all strains in the species. Previous pan-genome analysis of *Salmonella* has identified unique genetic landscapes of different serovars and strains and demonstrated the potential to understand strain relationships and phenotype differences^5–7^.

Recently, an independent component analysis (ICA)-based method has been successful in elucidating quantitative bacterial TRNs^8–10^. ICA is a signal separation algorithm that deconvolutes mixed signals into their individual sources and determines their relative strengths^11^. By applying ICA to bacterial transcriptomes, we can identify independently modulated sets of genes, called iModulons (individual source signals) and the activity level of the iModulon across different conditions (relative signal strength). Unlike regulons that are defined from the bottom up by experimental data of transcription factors and DNA binding, iModulons are derived computationally from gene expression via a top-down approach. It has been applied to transcriptomic datasets of *E. coli*, *B. subtilis* and *S. aureus* and provided valuable insights into the global TRNs of these species^8–10^.

In this study, we compute and analyze the iModulons of 6 Salmonella typhimurium strains, and compare their structure and activity across the serovar. We generate a large transcriptomic compendium by downloading all the publicly available RNA-Seq data of this serovar from NCBI Sequence Read Archive (SRA) and compiled expression profiles of 533 high quality samples. The large size and diversity of conditions in this dataset make it a valuable starting point for machine learning of the TRN. In order to understand the serovar as a whole and simultaneously characterize individual strains, we performed a pan-genomic analysis on 500 Typhimurium strains and defined the Typhimurium core genome with 172 strains. We performed ICA on both the core genome and the individual strain genomes and obtained over 400 robust iModulons in total. Many of these iModulons are highly consistent with known regulons, while others offer guidance for new discoveries.

## Results

### Compiling the *Salmonella* Typhimurium RNA-seq Compendium

To compile the RNA-seq compendium, we scraped the NCBI Sequence Read Archive (SRA)^12^ for all publicly-available *Salmonella enterica* RNA-seq data as of August 20, 2020. The strains and serovars were labeled using SRA metadata, when available, or linked literature (Supplementary Figure 1). In total, the initial compendium contained 1,444 expression profiles across 17 serovars. Within serovar Typhimurium, there were 1,174 expression profiles across eight strains.

Each expression profile was processed using a standardized pipeline^13^. After low-quality samples were discarded (See **Methods**), the final Typhimurium compendium contained 533 expression profiles from 46 BioProjects distributed across 6 Typhimurium strains **(Figure 1a**) (Supplementary File).

**Figure 1.**
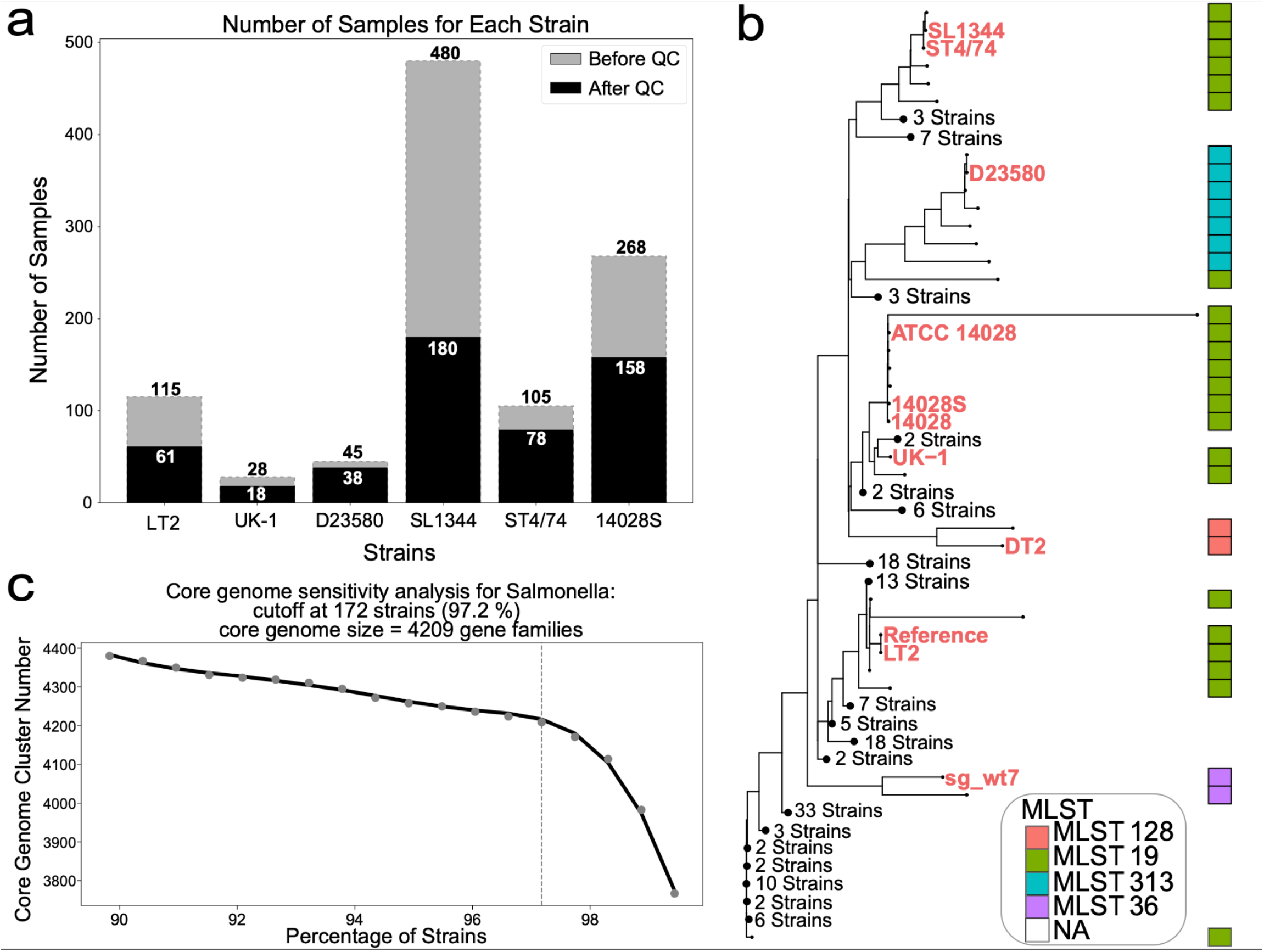
Dataset overview, sensitivity analysis and the phylogenetic tree. **a.** The number of samples for each strain before and after the quality control (QC) pipeline. This bar chart shows the distribution of available RNA-Seq data across 6 *S. enterica* Typhimurium strains. While SL1344 has the most number of strains, many failed QC because of the low number of reads mapped to coding sequences. The detailed quality control metrics can be found in **Methods**. **b.** Phylogenetic tree of 177 representative Typhimurium strains. Strains with names labeled in colors are strains with RNA-Seq data in our compendium. Strains DT2 and sg_wt7 each had only 4 samples. Due to the small size of the datasets, these two strains were used to build the core genome, but not included for individual ICA analysis. (See **Methods** section for details) The multilocus sequence typing (MLST) information of each strain (gathered from PATRIC) is presented in the bar on the right. The complete tree with uncollapsed branches can be found in (Supplementary Figure 6). Also note that strains ATCC 14028, 14028S and 14028 are considered as the same strain in subsequent analysis (See Supplementary Note 1). **c.**Sensitivity analysis for core genome definition. Many genomes from PATRIC are incomplete. To not lose too many gene families due to incomplete genomes, a sensitivity analysis was performed and a cutoff at 97.2% was chosen. Out of 177 genomes, 172 were used to define the core genome, resulting in a final core genome size of 4209 gene families.

### Pan-genomic Analysis Reveals the Core Typhimurium Genome

To understand the genetic diversity of serovar Typhimurium, we collected 3,329 *S. enterica* Typhimurium genomes from PATRIC^14^. After quality control (See **Methods**), we used CD-HIT^15^ to assemble the pan-genome matrix of 506 strains that includes all 8 strains we have RNA-Seq data for. The pan-genome contains 17,243 gene families. K-means clustering and explained variance were used to decide the optimal number of representative strains (See **Methods**). In the end, we identified 177 representative strains from which we generated the phylogenetic tree presented in **Figure 1b**.

Phylogenetic analysis of the pan-genome reveals the majority of the 177 strains to be ST19 strains, a major sequence type of S. enterica Typhimurium^16^. It is not surprising that 5 out of the 8 strains in our compendium are ST19 strains. The tree also shows that some of the RNA-Seq strains are more closely related than the others (SL1344 and ST4/74). The ST313 strain D23580 is a derivative from the ST19 strain ST4/74. Despite the small genetic difference of only 846 nucleotides, the two strains differ significantly in transcriptional signature and phenotypic features4. The phylogenetic tree clearly presents the relationship between the two strains, and guides our subsequent ICA analysis for strain-specific datasets. The pangenome defined in this study used 177 strains from S. enterica serovar Typhimurium. While smaller in scale and diversity as compared to previously defined Salmonella pangenomes^5–7,17^, this focused study is used to demonstrate the interoperability of pangenomic analytics and ICA. As we show later in the results, the combination of these tools guides our understanding of strain differences. Moreover, we want to use the phylogenetic tree to present the strains that we currently have RNA-seq data for and the diversity they represent within serovar Typhimurium.

To eliminate the negative effect of incomplete genome on the core genome size, a sensitivity analysis was performed on the 177 strains. With the cut off of 97.2%, 172 strains are used to define a core genome that contains 4,209 gene families (**Figure 1c**). For subsequent analysis of the core genome, we used a core compendium containing all 533 expression profiles, keeping only the 3,886 core genes (See **Methods**).

### Independent Component Analysis Captures Transcriptional Regulatory Network of *S. enterica* Typhimurium

To infer the TRN of *Salmonella* Typhimurium, we applied ICA to the core Typhimurium RNA-seq compendium, resulting in 115 robust iModulons. Together, these 115 iModulons explain 75% of the variance in gene expression (Supplementary Figure 2). Many statistical methods use explained variance simply to measure the quality of the reconstruction mathematically (e.g. PCA). However, ICA results carry biological meanings and upon characterization, the independent components can be linked to transcriptional regulations. Therefore, the explained variance of ICA offers not only mathematical representations but also biologically-relevant explanations of variance in gene expression.

To characterize the 115 iModulons, we first constructed a draft TRN of *S. enterica* Typhimurium using a pre-defined TRN for the closely related bacteria *Escherichia coli*^8^, a model organism which has a more well-characterized TRN. We subsequently compared each iModulon against each known regulon and found that 60 iModulons had significant overlap with known regulons (See **Methods**). These “Regulatory” iModulons recapitulate the structure of known regulons, and demonstrate that ICA of one model species can be highly informative for characterization of closely related species.

The relationship between Regulatory iModulons and known regulons can be measured using two metrics: iModulon recall, which is the fraction of iModulon genes that are shared by the regulon, and regulon recall, which is the fraction of regulon genes that are shared by the iModulon. The scatter plot in **Figure 2a** shows the distribution of Regulatory iModulons in terms of their concordance with a known regulon. The histograms on the sides show high recall rates for many of the regulatory iModulons, demonstrating good agreement between the iModulon TRN and the previous TRN structure.

**Figure 2.**
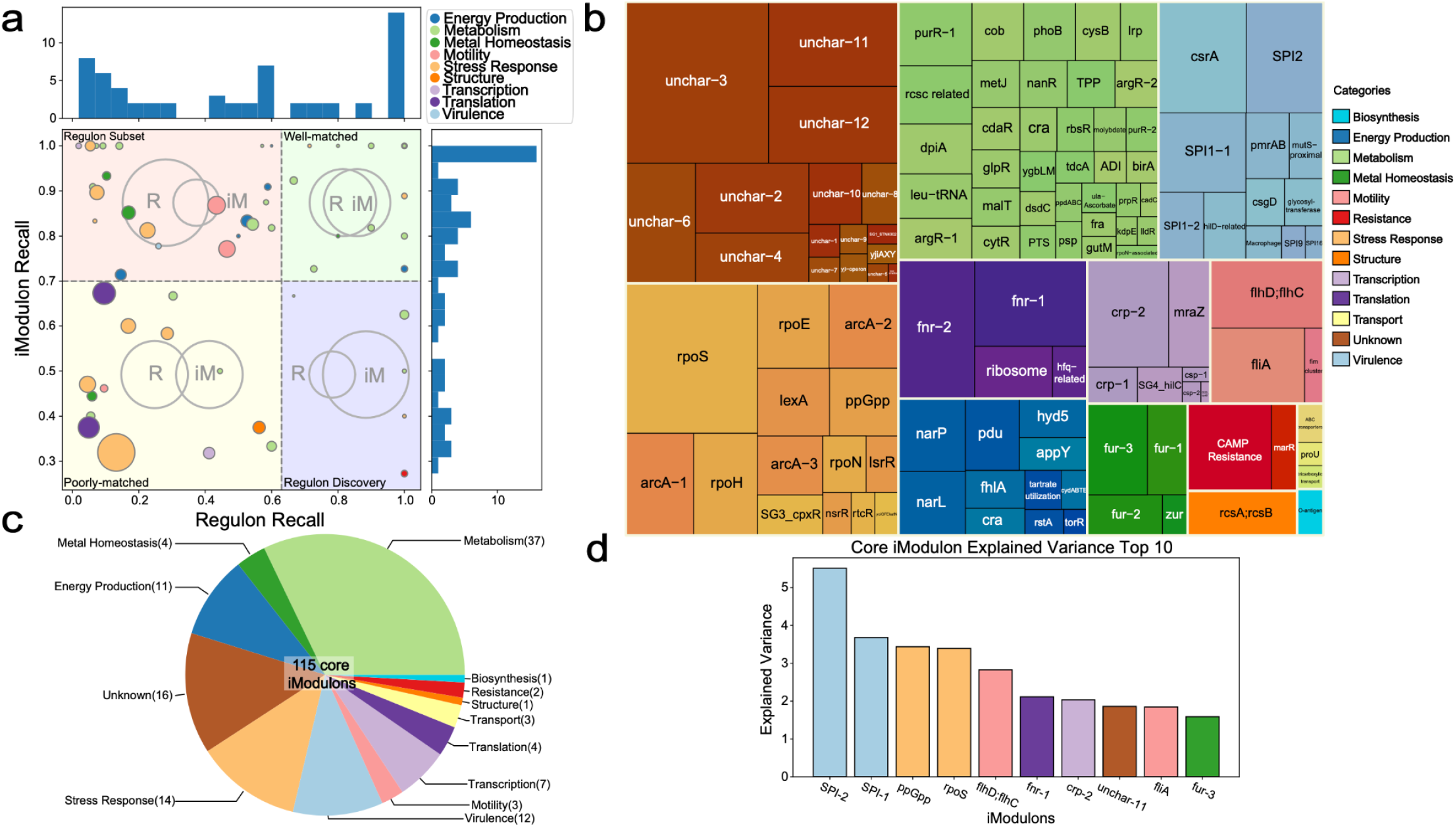
Core iModulon Statistics. **a.** Scatter plot of all the regulatory iModulons with histograms on the side. Each dot represents a regulatory iModulon. The size of the dot is determined by the size of the iModulon. Regulon recall is defined as (# of shared genes between iModulons and regulon) / (# of genes in a regulon). iModulon recall is defined as (# of shared genes between iModulons and regulon) / (# of genes in an iModulon). High iModulon recall and regulon recall values indicate high consistency of the iModulon with a previously characterized regulon. The regulatory iModulons are divided into four quadrants: Regulon Subset (top left), Well-matched (Top right), Poorly-matched (bottom left) and Regulon Discovery (bottom right). The relationship between the regulon (R) and iModulon (iM) in each quadrant is shown by the Venn diagram in the background. **b.** Core iModulons and categories. The names and functional categories of 115 core iModulons are presented in this tree map. The box sizes represent the size of the iModulons. The number of iModulons in each category can be found in the category breakdown pie chart in panel **c**. **d.** Ten iModulons with the highest explained variance. iModulons with important functions such as virulence, stress response, global regulatory network and motility were captured.

Information from public databases such as Gene Ontology (GO) enrichments and KEGG PATHWAY Databases were used to characterize the remaining 55 iModulons. These iModulons represent genes with coordinated actions but no known regulon. Some of them are from related biological pathways that are regulated together to achieve a biological function, while some less characterized gene sets can be targets for regulon discovery. To describe these iModulons, we define Genomic iModulons as iModulons that account for changes to the genome such as single gene knock-outs, large deletions, and or duplications of genomic regions. Functional iModulons are defined as iModulons that contain genes enriched for a specific function, but are not linked to a specific transcriptional regulator^18^.

To further evaluate the scope of the iModulon structure, we assigned each iModulon to a functional category (**Figure 2b,c**). Many iModulons were related to metabolism (which include carbohydrates, amino acids, nucleotides and lipids metabolism), stress response (cold/heat response, osmotic stress response, anaerobic shock) and energy production. It is also interesting to see iModulons that relate to virulence and resistance, which are essential mechanisms for understanding and combating *Salmonella* infections. In fact, the two iModulons that capture the highest fraction of expression variance are related to the *Salmonella* pathogenicity islands (SPI-2 and SPI-1, respectively). Other iModulons that explain large fractions of expression variance include stress response iModulons (ppGpp and RpoS), and motility iModulons (flhD;flhC), indicating the importance of these mechanisms (**Figure 2d**). Uncharacterized iModulons with high explained variances (e.g. unchar-11) can represent important biological functions and represent good targets for further studies.

### ICA extracts *Salmonella* pathogenicity islands

Many of the virulence genes of *Salmonella* are located in *Salmonella* pathogenicity islands (SPIs)^19^. SPIs contain important genes that *Salmonella* needs to invade, survive and spread in the host. There are 12 common SPIs in *Salmonella* Typhimurium^20^. ICA extracted 5 iModulons related to 4 SPIs (SPI-1, SPI-2, SPI-9 and SPI-16) in the core genome. The SPI-1 and SPI-2 iModulons are the top 2 iModulons with the highest explained variance (**Figure 2d**).

SPI-1 encodes a type three secretion system (T3SS) that delivers effector proteins to help *Salmonella* penetrate the intestinal epithelium. There are 46 genes located in SPI-1^21^, which includes genes coding for the secretion system apparatus, effector proteins, and their regulators. ICA extracted 2 iModulons enriched for SPI-1 genes, in total accounting for 5.2% of the global expression variance across the compendium. The largest SPI-1 iModulon contains not only genes from SPI-1, but also four genes from SPI-4. (**Figure 3a**). It is known that both SPI-1 and SPI-4 are required for *Salmonella*’s entry into polarized epithelial cells^22^, and studies have shown that transcriptional factors encoded in SPI-1 such as *hilA*, *hilC*, *hilD*, and *sprB* have a regulatory effect on SPI-4 genes^23,24^. *hilD* resides in SPI-1 and codes for the important virulence regulator HilD. HilD regulates many virulence genes in SPI-1 directly and interacts with other virulence-related regulators including HilA, HilC and RtsA (outside of SPI-1). HilD forms a feedforward loop with HilC and RtsA to amplify the regulation on HilA, which is known to regulate sprB and genes in SPI-4^23,25^. Not surprisingly, in the Differential iModulon Activity (DIMA) plot in **Figure 3b**, iModulons related to *hilC*, *hilD* and *sprB* have lower activities along with the two SPI-1 iModulons in the *hilD* mutant samples (PRJNA315446). Interestingly, when focusing on *hilC*, *hilA, sprB* or *rtsA* mutants, we notice that the activities of the SPI-1 iModulons did not vary much compared to the wild type.Besides an iModulon directly related to the gene that was knocked out, we do not see iModulons of related regulators with differential activities. (Supplementary Figure 3). This confirms the significance of HilD in the regulation of SPI-1. It is also consistent with the previous findings that HilD has a more dominant role compared to HilC and rtsA^25,26^.

### The CRP iModulon elucidates the structure of the Typhimurium CRP network and its effect on antibiotic susceptibility

The cyclic AMP Receptor Protein (CRP) is a global regulator that orchestrates a variety of biological pathways. In *E. coli*, CRP is known to control 70 genes for transcription factors, affecting over 300 gene targets involved in metabolism and stress response^27^. Not surprisingly, in *S.enterica* Typhimurium, CRP also carries important roles, and deletion of the crp gene significantly affects virulence and metabolism^28^. The complete CRP network of *Salmonella* is yet to be defined, and the *crp* iModulon provides a potential structure of the Typhimurium CRP network and offers guidance to find targets of CRP. This *crp* iModulon contains 43 genes, 16 of which are known to be regulated by CRP. A large number of the genes in the iModulon are transcription factors that subsequently regulate more biological activities (**Figure 3c**). Genes crucial for metabolism are also a significant component of the iModulon, which agrees with our predictions from the *E. coli* network. There are 14 uncharacterized genes in the iModulon that make good targets for further studies.

**Figure 3.**
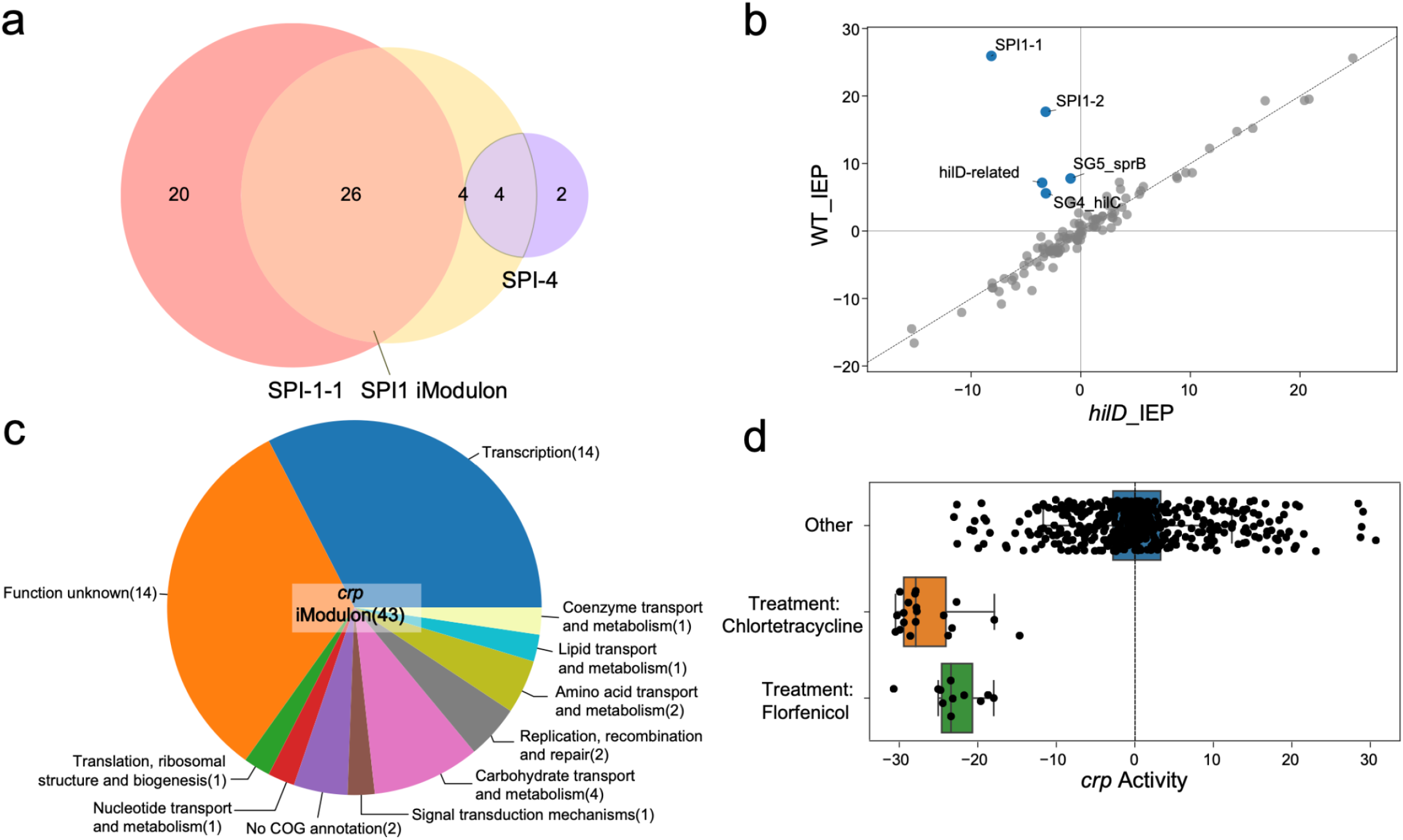
iModulons validate previous results and provide insights on virulence and the crp regulatory network. **a.** Relationship between the SPI-1, SPI-4 pathogenicity islands and the SPI-1-1 iModulon (the larger SPI-1 iModulon). The SPI-1-1 iModulon contains 4 genes in SPI-4, 26 genes in SPI-1 and 4 genes in neither. The four genes are STM1328, STM4312, STM4313 and STM4315. **b.** Differential iModulon Activity (DIMA) plot for wildtype samples and *hilD* knockout samples in intermediate exponential phase (IEP) (PRJNA315446). iModulons related to SPI-1, SPI-4 and the coordination between these two islands are found to have differential activity levels between wildtype and *hilD* knockout samples (PRJNA315446). **c.** Gene function category breakdown of the *crp* iModulon. The numbers in parentheses give the number of genes in each category. **d.** The activity level of the crp iModulon under the treatments of chlortetracycline and florfenicol. The activity of this iModulon significantly decreases under both conditions.

The activity levels of each iModulon under different conditions indicate the activity of the underlying biological signal and regulator. The activity of the *crp* iModulon shows that it is strongly repressed under two conditions with antibiotic treatments (PRJNA344670) (**Figure 3d**). The activity of the iModulon drastically dropped when the multidrug resistant cells are treated with chlortetracycline and florfenicol, antibiotics that are frequently administered to animals for disease prevention. It is known that deletion of *crp* increase *Salmonella*’s resistance of fluoroquinolone antibiotics, possibly by affecting the drug delivery system and altering the drug target with DNA supercoiling^29^, but the change in expression of the *crp* regulon under the treatment of tetracyclines or florfenicol is undocumented. Unlike fluoroquinolone which inhibits DNA topoisomerases, both chlortetracycline and florfenicol work by interacting with the bacterial ribosome to prevent peptide synthesis. It is possible that downregulation of the *crp* iModulon in MDR samples results from the regulation of drug deliveries similar to the mechanism to resist fluoroquinolone. An alternative hypothesis is that among the targets of *crp*, some ribosomal protein genes or other translation-related genes can influence the antibiotic’s interactions with the ribosome. Thus, the repression of the *crp* regulon could be part of the response mechanism of the multidrug resistant strains to reduce their susceptibility to antibiotics.

### Three transport iModulons are activated by to antibiotic stress in MDR strains

The emergence of multidrug resistant (MDR) strains poses a serious challenge in preventative care and infection treatment. To help understand how these strains adapt to antimicrobial agents, we investigated three iModulons (the proU iModulon, the molybdate iModulon, and the NikR iModulon) that have high activities under antibiotic treatment of MDR strains in PRJNA344670 (Supplementary Figure 4).

The proU iModulon (*proXWV*) encodes an osmoprotectant transport system activated by osmotic stress. Unexpectedly, when the MDR strains are treated with subinhibitory concentrations of antimicrobials agents, the activity of this iModulon drastically increases (Supplementary Figure 4). It has not been documented that the proXWV operon is associated with other cellular functions in *Salmonella*, but the results from ICA indicate that the activation of this transporter can be triggered by antimicrobial agents.

The molybdate iModulon consists of 7 genes (*modA*, *modB*, *modC*, STM0770, STM0771, STM3142 and *yghW*), many of which are a part of the essential molybdate transport system. This system delivers molybdate oxyanions into the cell, where they are converted to molybdenum (Mo) and serve as cofactors for many enzymes. While it is possible that more Mo is required for molybdoenzymes involved in the drug resistance mechanisms, the high activity for this iModulon in MDR strains implies that molybdate compounds affect bacterial survival under antibiotic treatment. Evidence has shown that silver molybdate and copper molybdate can enhance the abilities of antimicrobial agents in killing other Gram-negative bacteria such as *E. coli*. The metal ions can disrupt the cell membranes while the molybdenum oxide or molybdate oxyanion modulates the local pH to inhibit further growth^30^. It is still unclear why Typhimurium MDR strains upregulate molybdate transport genes when treated with antibiotics, but this result strengthens the argument that molybdate has a major effect on the survival of Gram-negative bacteria, especially when antimicrobial agents are present.

Finally, the NikR iModulon consists of five putative ABC transporter genes in the NikR regulon (STM1255 - STM1259). Not only is this regulon important for the regulation of nickel uptake, all 5 genes are found in the quorum sensing pathways as well^31^. Previous studies have found that nickel chelator reduces the virulence and survival rate of multiple enterobacteriaceae including *S. enterica* Typhimurium^32^ by sequestration. The enhanced activity of the NikR iModulon under antibiotic treatments reinforces the hypothesis that nickel plays an important role in the survival of MDR Typhimurium strains and invites explorations for nickel-related methods to combat *Salmonella* infections.

### Uncharacterized iModulons provide future directions for investigation

Many iModulons can be characterized with existing knowledge, but some others are not previously documented in literature, giving us new directions for future research. One iModulon (uncharacterized-1) contains five genes that seem to relate to PTS sorbitol transporter (**Figure 4a**). These five genes may reveal the structure of an new operon, with STM2749 as the putative regulatory element. The potentially new operon shows increased expression for *csrA* mutant stains, suggesting that it is repressed by this global regulator. Furthermore, the low activity of the iModulon for *csgD*-perturbed cells the regulatory role of *csgD* on these genes (**Figure 4b**). The DIMA plot in **Figure 4c** also shows the close relationship of this iModulon and the *csgD* iModulon, as they are the only two iModulons with lower activities in *csgD* mutant samples. *csrA* regulates virulence, metabolism, and biofilm formation, while *csgD* is a central biofilm regulator in *Salmonella*. Therefore, we hypothesize that this gene cluster may be related to biofilm formation.

**Figure 4.**
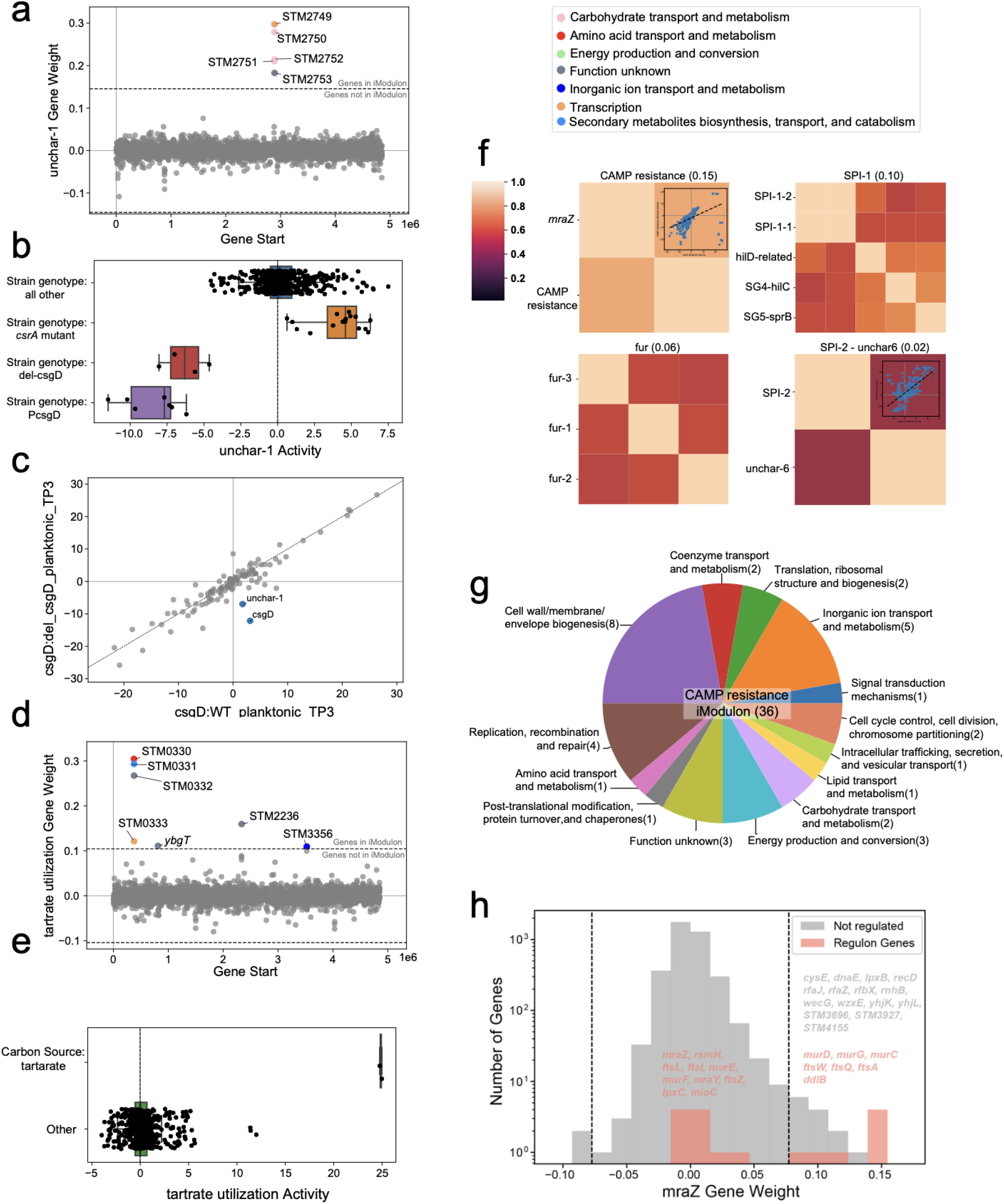
iModulons guide new discoveries. **a.** iModulon composition and gene weights graph of iModulon uncharacterized-1. Five genes are in the iModulon, with one potential transcriptional regulator. **b.** Activity of uncharacterized-1 iModulon for *csrA* and *csgD* mutant samples. It can be seen that the iModulon has high activity when *csrA* is knocked out (PRJNA421560), or when *csgD* is disturbed (PRJNA280002^33^). **c.** Differential iModulon activity plot (DIMA plot) of a *csgD* KO sample and a wildtype sample in planktonic culture. **d.** iModulon composition and gene weights graph of tartrate utilization iModulon. The activity of this iModulon is shown in panel **e**, where we see increased iModulon activity when the samples are treated with the single carbon source tartrate (PRJEB4981^34^), and when the strains carry a mutation in the Rho factor (PRJEB34015). **f.** Top four iModulon activity clusters with the highest correlations. **g.** Gene function category breakdown of the CAMP resistance iModulon. This pie chart shows the number of genes in each functional category in the CAMP resistance iModulon. **h.** Histogram of gene weights for the *mraZ* iModulon and regulon. The two dashed lines indicate the cutoff gene weight values for defining the iModulon. Genes in the mraZ regulon are indicated in red. Seven genes in the *mraZ* regulon is captured by the iModulon.

In another case, the tartrate utilization iModulon contains genes with a variety of different functions, but are not known to be regulated together (**Figure 4d**). Interestingly, this iModulon is found to have increased activity when tartrate is the only carbon source (**Figure 4e**) (See Supplementary Note 2). Among the 7 genes in the iModulon, STM3356 is a putative cation transporter whose function is still unknown, but disruption of this ORF can convert tartrate fermenting phenotypes into tartrate non-fermenting phenotypes for *S. enterica* serovar Paratyphi B dT+^35^. The presence of this gene in the *S. enterica* Typhimurium core genome ICA results suggests that this gene is crucial for the utilization of tartrate for Typhimurium as well. For the rest of the genes in this iModulon, STM0330 is a 3-isopropylmalate dehydratase, and STM0331 and STM0332 are putative hydrolases. STM0333 is a putative LysR family transcriptional regulator and ybgT is the subunit of Cytochrome bd-I oxidase. None of the genes are documented to be directly related to the conversion of tartrate.

### iModulon clusters reveals coordinated activities and illuminate CAMP resistance mechanism

The activity fluctuation of individual iModulons under specific conditions can help us understand the functions of one set of genes. However, biological processes often involve multiple gene sets with coordinated functions. These genes are not necessarily regulated by the same regulator but they all respond together to the same stimulus, forming a stimulon. These gene sets can be reflected by iModulons that are activated together under specific conditions and have similar activities across the entire compendium. To investigate the relationships between iModulons and the biological insights of these relationships, we clustered the iModulons based on their activities. iModulons in the same cluster are likely to have related biological functions, since their activities are correlated. This is indeed the case when we further examine clusters with the highest correlations: one cluster with the high correlation score contains five iModulons directly related to the SPI-1 pathogenicity island. The *fur* cluster consists of three iModulons related to the ferric uptake regulator Fur.(**Figure 4f**).

The clear relationships between iModulons in the SPI-1 and *fur* clusters give us enough reason to believe that iModulon activity clustering is a reliable way to infer iModulon relationships and derive biological understanding. Even though in some other clusters the connection can be harder to decipher, we believe that they do represent true biological implications. One cluster that is worth mentioning is the cationic antimicrobial peptide (CAMP) resistance cluster that sheds light on Typhimurium’s defense mechanism against CAMPs. CAMPs are microbicidal peptides produced by the immune system of many organisms to combat microbial infection. *Salmonella* and other gram-negative bacteria are major targets of CAMPs, so the bacteria have developed mechanisms to protect themselves. One of the most extensively studied resistance mechanisms is surface remodeling. Since the outer surface is the first line of defense against CAMPs, *Salmonella* has developed ways to enhance resistance through modifying its surface. These include reducing the negative charges on the surface to limit electrostatic interactions with CAMPs, regulating O-antigen length to create a stronger barrier, decreasing membrane fluidity to control CAMPs intake, and increasing efflux of CAMPs^36^.

In the CAMP resistance iModulon activity cluster, we captured 2 iModulons that potentially act in concert to resist antimicrobial peptides. The CAMP resistance iModulon contains 36 genes, 5 of which (*pqaB*, *sapD*, *sapF*, STM2300, STM2303) are in the CAMP resistance pathway from the KEGG PATHWAY Database (stm01503). The genes in this iModulon have a variety of functions including cell wall and membrane biosynthesis (**Figure 4g**). The *mraZ* iModulon consists of 22 genes, 7 of which are regulated by *mraZ*, a gene within the *dcw* (division and cell wall) cluster which has a major effect on cell division and peptidoglycan synthesis (**Figure 4h**). Many genes in the two iModulons are related to biosynthesis and transport of lipopolysaccharide (LPS). This result aligns well with the LPS modification hypothesis, where *Salmonella* needs to redesign its LPS to be less anionic to evade interactions with CAMPs. Genes related to O-antigen biosynthesis are present in both iModulons, pointing to the regulation of O-antigen length. We also found genes related to peptidoglycan (PG) synthesis in this iModulon cluster. There is little evidence of the direct involvement of PG remodeling in *Salmonella*’s defense response to CAMPs. However, PG is closely enveloped by LPS, so it is also possible that LPS remodeling works jointly with PG to protect the inner membrane from damage. Other than genes related to synthesis of the cell surface components, the presence of several transporters can be the indication of flux and binding regulation at the cell membrane. Moreover, we also discovered several genes important for DNA replication, repair, and recombination. Antimicrobial agents such as CAMPs can induce DNA damage directly and indirectly, so these genes are likely part of the SOS response of the bacteria to repair and synthesize more DNA for survival. The rest of the genes with unknown functions are likely to be involved in the response of CAMPs, and this iModulon cluster offers insights into future investigations of antimicrobial resistance.

While SPI-1 is crucial for epithelial cell invasion, SPI-2 is required for intracellular replication, survival and persistence. The SPI-2 iModulon explains 5.5% of the compendium-wide expression variance. It is clustered with an uncharacterized iModulon **(Figure 4f)**. The uncharacterized-6 iModulon contains many virulence and resistance-related genes including *pagP*, *utgL*, the PhoP/Q two-component regulatory systems and genes this system regulates. These genes all show close relationships with the SPI-2 island. However, the relationship between virulence and many other genes are less obvious, and around half of the genes are characterized to code for putative proteins with unclear functions. These poorly characterized genes warrant further study to explore their relationship with pathogenicity islands and virulence.

### Comparative studies guides investigation of stress response in related strains

One of the projects that we looked at was designed to understand the global gene expression differences between D23580 (ST313 isolate) and its ancestral strain ST4/74 (ST19 isolate) under infection-related conditions (PRJNA490148)^4^. The genomes of these two strains are 95% identical, but each strain has distinct phenotypic characteristics. Identifying the genes whose expression profile varies in the two strains may assist in explaining the phenotypic differences. Samples from both strains were cultured under the same infection-related conditions. This comparative study allows us to compare iModulon activities for the same conditions in different strains, and we discovered three uncharacterized iModulons that displayed interesting activity patterns.

The iModulon Uncharacterized-5 consists of only two genes: *yaaY*, an uncharacterized gene and *yccX* that codes for acylphosphatase. The Uncharacterized-12 iModulon is quite large and contains 49 genes. While there are genes related to virulence (*ssrA/ssr*B two component system) and fimbriae (*fimY* and *safB*), the majority of the genes in this iModulon code for putative proteins spanning a variety of functional categories (**Figure 5a**). Although the functions of these two iModulons still remain obscure, their activities strongly indicate that they are related to ST4/74-specific mechanisms. In **Figures 5b** and **5c**, the activities of the two iModulons were shown across all 6 strains. For samples from all the other strains, this iModulon has an activity around zero. However, for strain ST4/74, the activities for both iModulons range from −20 to 10. The small cluster of samples with low activities in ST4/74 are in a variety of conditions, so it is unclear what triggers the drop of iModulon activities.

**Figure 5.**
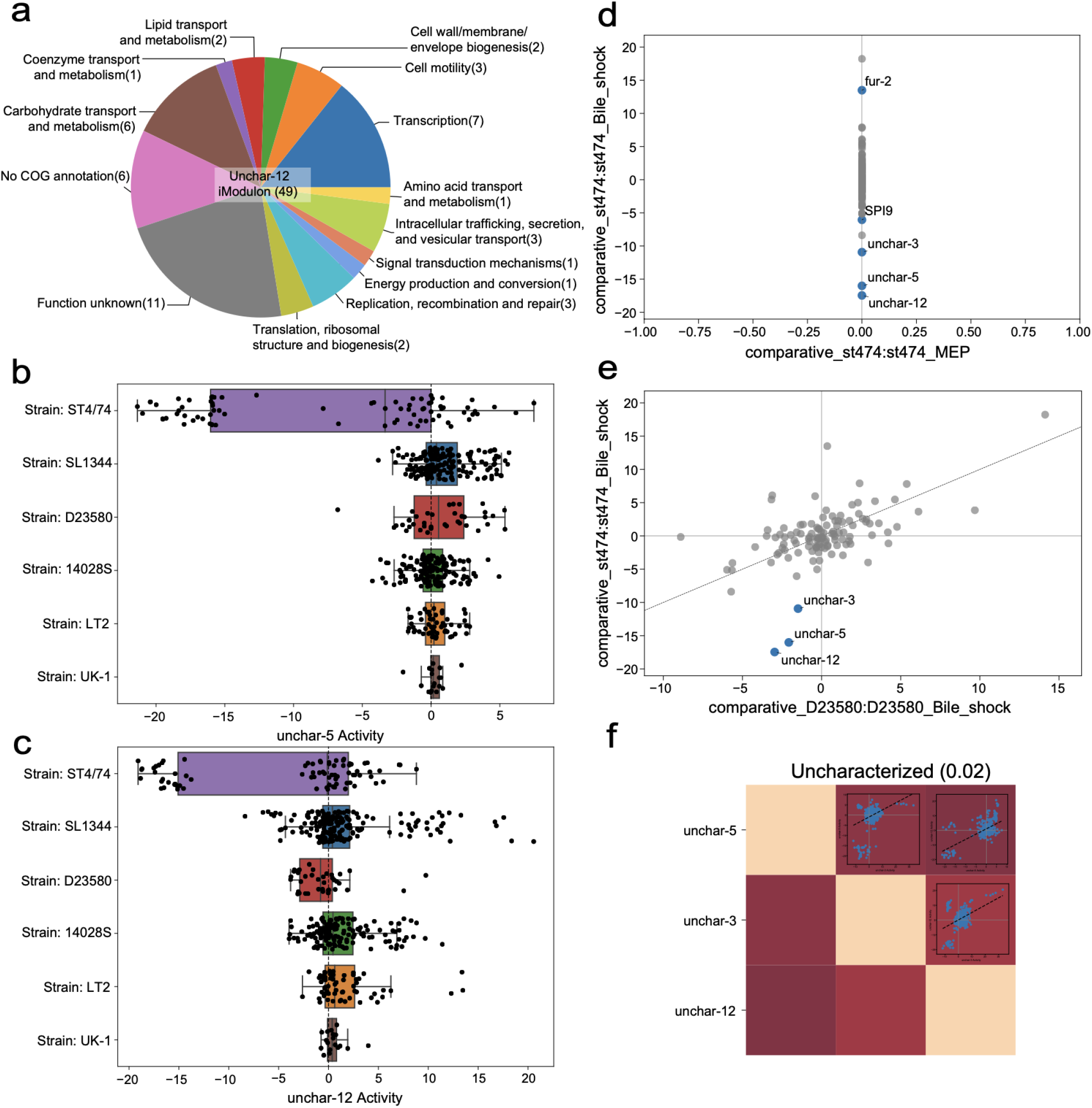
Comparative studies reveal iModulons that potentially contribute to bile response in strain ST4/74. **a.** Gene function category breakdown of the uncharacterized-12 iModulon. **b.** The activity of iModulon uncharacterized-5 across samples from all six strains. While for five strains the activity of this iModulon is close to zero, for strain ST4/74, several samples showed significantly decreased activity of this iModulon (these samples are from PRJNA215033, PRJNA315446, PRJNA393682, PRJNA490148). No obvious patterns were seen in these samples with low activities. **c.** The activity of iModulon uncharacterized-12 across samples from all six strains. Samples from strain that showed significantly decreased activity are from PRJNA215033, PRJNA315446, PRJNA393682, PRJNA490148). **d.** Differential iModulon Activity (DIMA) plot for strain ST4/74 samples under bile shock against samples in LB at middle exponential phase. **e.** Differential iModulon activity plot for samples under bile shock in strain ST4/74 and D23580 **f.** iModulon activity clusters for three uncharacterized iModulons that show low activity for samples under bile shock in strain ST4/74.

However, looking at the DIMA plots of the bile shock samples from strain ST4/74 compared to ST4/74 samples cultured in LB and bile shock samples from strain D23580 (PRJNA490148), we speculate that these two iModulons play a role in the bile response for ST4/74. (**Figures 5d, e**). It can be seen that the uncharacterized-5 and uncharacterized-12, along with iModulon uncharacterized-3 show low activities under bile shock in strain ST4/74. Uncharacterized-3 is a large iModulon (113 genes) with no enrichment in any previously-defined regulon or pathways, but the functional category breakdown of the gene content can be found in Supplementary Figure 5. The activities of these three iModulons are also found to be correlated as shown in **Figure 5f**. To our knowledge, strain ST4/74 is not particularly sensitive or resistant to bile salts compared to D23580 or other strains in this study. However, this strain was initially isolated from the bowel of a calf with Salmonellosis, so it is possible that it utilizes a special set of genes to interact with bile salts in the bowel. The exact mechanism remains to be determined, but the ICA results offer guidance to further explorations of this strain-specific response. More ICA results related to the comparisons of these two strains can be reproduced with the materials at https://github.com/AnnieYuan21/modulome-Salmonella.

### Composition of pathogenicity island iModulons differ in content across the strains

iModulons from the core genome can inform us about the TRN of serovar Typhimurium as a whole. However, different strains within the serovar have different properties, and these properties can be explored using strain iModulons. Here we examine how evolutionary pressures shape both the genome composition and the transcriptional regulatory network of individual strains.We generated six individual strain-specific RNA-seq datasets and ran ICA on them, obtaining six sets of strain iModulons. The number of iModulons and their category breakdown are presented in **Figure 6a**. With both the core and all the strain iModulons, we were able to do some comparisons.

**Figure 6.**
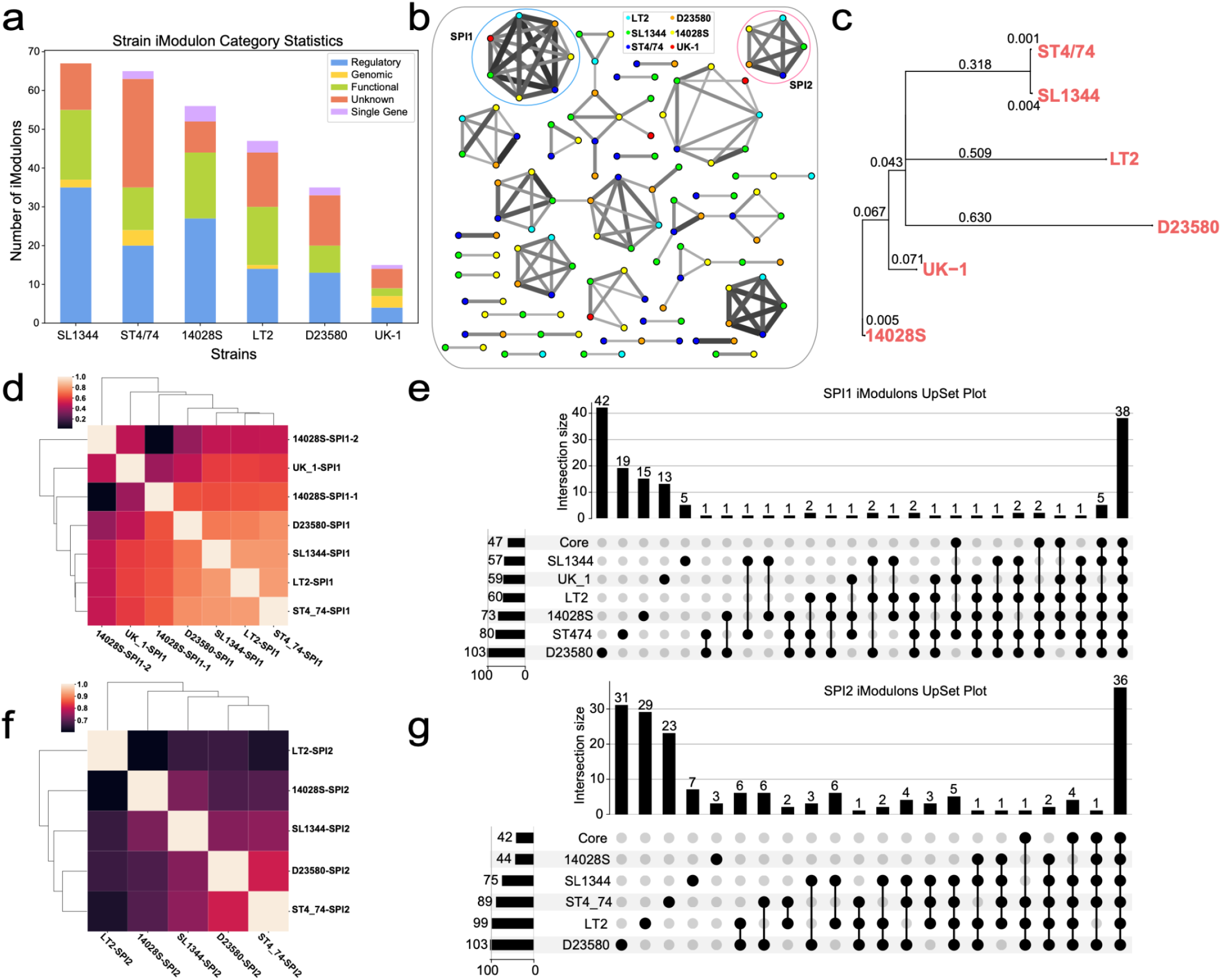
iModulon comparisons in different strains. **a.** The number and category of iModulons for all six strains. **b.** Reciprocal Best Hit (RBH) graph of iModulons from 6 strain datasets. Each node is an iModulon and each edge is the RBH. The color of the node corresponds to the strains and the thickness of the edge indicates gene weighting similarity. The cutoff chosen in generating this RBH graph is a Pearson R value of 0.3^53^. Two clusters that are circled indicate the SPI-1 and SPI-2 iModulons. The full labeled RBH graph can be found in Supplementary Figure 7. **c.** Simplified phylogenetic tree with only the 6 strains used for ICA analysis. The branch lengths are indicated by the scores. **d.** Cluster map for all SPI-1 iModulons in the 6 strains. **e.** Upset plot to compare the gene contents of the SPI-1 iModulons across the core genome and the strains. **f.** Cluster map for all SPI-2 iModulons in the 5 strains (UK-1 does not have a SPI-2 iModulon). **g.** Upset plot to compare the gene contents of the SPI-2 iModulons across the core genome and 5 strains.

If iModulons with similar composition and function are present in more than one strain, investigating the differences in iModulon composition for each strain can help understand strain-specific genetic signatures that account for differences in phenotypic features. The RBH graph in **Figure 6b** shows clusters of similar iModulons in the strains. Here we use the SPI iModulons as an example. Four SPIs were identified in the core genome, with SPI-1 and SPI-2 having the highest explained variance. Both of these two islands are very important for the virulence of Typhimurium, and showed up in the strain iModulons as well. SPI-1 were found in all six of the strains, while SPI-2 were found in five. Note that strain UK-1 does not have an SPI-2 iModulon. Since UK-1 does have an SPI-2 island, this observation can simply be the result of the relatively small RNA-seq dataset that UK-1 has.

To help understand the iModulon differences, the phylogeny of the 6 strains are presented in a simplified phylogenetic tree in **Figure 6c** with the phylogenetic distances labeled on the branches. The clustermap of all the SPI-1 iModulons in **Figure 6d** shows that the SPI-1 iModulons across all the strains are overall correlated. Interestingly, strains such as LT2 (avirulent) and D23580 (ST313) are clustered with the two closely related ST19 strains ST4/74 and SL1344. Also, the two SPI-1 iModulons in strain 14028S have poor correlation with each other. The larger SPI-1 iModulon codes for the majority of the SPI-1 genes with various functions including the regulators, invasion factors and T3SS apparatus, while the smaller one codes mostly for effectors and invasion factors. While most genes unique to the 14028S SPI-1 iModulons are uncharacterized, one effector GtgA was identified that is specific to this strain.

The differences in the SPI-1 iModulon contents are shown by the upset plot in **Figure 6e**. The strains and the core share 38 genes in the SPI-1 iModulons, while many strains also have their unique gene sets related to the SPI-1 island. The most unique strain is D23580, with 42 unique genes in its SPI-1 iModulon. Strain D23580 is an extremely invasive ST313 strain associated with non-typhoidal gastroenteritis and systemic disease. A multiple-antibiotic resistant regulatory gene, *marB*, was identified uniquely in D23580 SPI-1 iModulon, which supports its multidrug-resistant phenotype. The sequence of D23580 demonstrates genomic degradation, a hallmark for host adaptation. Many pseudogenes in this strain are homologous to genes in typhoidal serovars such as Typhi and Paratyphi, so it is hypothesized to be an intermediate between non-typhoidal serovars and typhoidal ones^37^. The unique genes in its SPI-1 iModulons are highly enriched for the KEGG flagellar assembly pathway (map02040) and the *flhDC* regulon. Flagella motility is extremely important for the infection of epithelial cells. Flagellar genes are under tight control of regulators coded in SPI-1 and they coordinate closely with the SPI-1 T3SS during infections^38,39^. The flagellar genes of D23580 are previously shown to be differentially expressed compared to ST19 strain ST4/74. It was also found that D23580 has attenuated expression of flagellin which allows it to cause minimal inflammatory response and reduce host cell death. This adaptation significantly enhances its survival within macrophages and resembles Typhoidal serovars that also infect the host cells while causing minimal inflammatory responses^37^. The unique genes in the D23580 SPI-1 iModulon further strengthens the argument that flagellar genes play a crucial role in virulence and inflammation, and accounts for the more invasive and resistant phenotype of D23580.

Compared to SPI-1, the SPI-2 iModulons are less correlated across the strains. (**Figure 6f**). Other than strains D23580 and ST4/74, the rest of the strains have low correlation scores for this iModulon. Nonetheless, all strains and the core still share 36 genes in the SPI-2 iModulon (**Figure 6g**). Strain D23580 again possesses the highest number of unique genes, five of which are part of the Vancomycin resistance pathway (map02020). These genes point to the multidrug resistant properties of this strain being tightly linked to virulence. Other interesting observations include that the LT2 SPI-2 iModulon contains 6 ABC transporters genes mainly the *malEGFK* operon and the *proVXW* operon operon. These LT2 specific SPI-2 related genes are also known to be regulated by *rpoD* (10 genes) and *rpoS* (3 genes). An defective *rpoS* gene is found to be the sole cause of avirulence of LT2^40^. The rest of the virulence mechanisms are assumed to be complete. It is unclear why these genes show a close relationship with the LT2 SPI-2, but investigations of these genes might lead to deeper understanding of the genomic roots of LT2’s unique phenotype.

### Strain iModulons captures prophage presence in different strains

Prophages can contribute to bacterial virulence and affect the phenotypic features of the bacteria. The prophage repertoire of *Salmonella* is diverse, and each strain tends to have its own unique set of prophages. ICA decomposition of the individual strain datasets extracted prophage-related iModulons that indicates the existence of certain prophages in each strain. For example, the Fels-1 and Fels-2 prophage-related iModulons are only found in LT2^41^. Hypothesis exists that these two phages are lost in the virulent strains^41^. Genes related to the most common prophages to *Salmonella* Typhimurium Gifsy-1 and Gifsy-2 prophages are found across all strains except UK-1 and D23580 and the latter inactivates these two prophages^42^. We also discovered a Gifsy-3 like iModulon in strain 14028S. This Gifsy-3 iModulon shows high correlation with the Gifsy-1 iModulon in strain SL1344. This might account for the similar immunity module of these two prophages^43^. Moreover, it was observed that the prophage iModulons in all strains except LT2 and D23580 carry part of the ST64B prophage. This phage is inactivated in strain D23580 but is completely absent in strain LT2. ST64B codes for the effector SseK3 and it was proposed that the presence of this phage enhances *Salmonella* survival in the blood^44,45^. Our ICA results are highly consistent with the documented existence of prophages in the studied strains. Moreover, a deeper examination of the iModulons can also reveal the difference in iModulon contents for the same prophage to help understand regulations of prophage genes and how they affect strain virulence.

## Discussion

*Salmonella enterica* is one of the leading causes of foodborne illnesses globally. While Typhimurium is one of the most studied non-typhoidal serovars, the phylogenetic tree in **Figure 1b** shows the limited number of strains that we currently have RNA-seq data and points to the amount of understanding we still need to develop for more strains in serovar Typhimurium. We have little information about the transcriptional signatures of the majority of the strains in this serovar and clades at the bottom of the tree are poorly understood. The available datasets cover mostly the ST19 group and many strains in this group are in close relationships with each other. Even though these strains are more common, they do not represent the transcriptomic landscape of the entire serovar. It might be worthwhile to inspect some less common strains from different branches. By comparing and contrasting expression profiles, we might be able to discover traces of strain evolution or genetic signatures that account for characteristics of different Typhimurium strains. It was demonstrated in this work that ICA and pan-genomic tools can guide the investigation of strain differences and relationships. If we experimentally validate the relationship between differential genetic features identified from ICA and different phenotypes, we can potentially predict phenotypes (such as pathogenicity, resistance and persistence) of rare strains and new clinical isolates to design treatment methods efficiently.

Moreover, non-typhoidal *Salmonella* is becoming an increasingly serious threat to public health with the rise in multidrug-resistant strains^46,47^. The excessive use of antibiotics in clinical settings and agricultural practices are increasing bacteria’s tolerance to drugs and lowering *Salmonella*’s susceptibility to common antibiotics, significantly reducing treatment success rates. The development of antibiotic resistance can be genetic or phenotypic, and varies across strains and serovars^48,49^. Additionally, The molecular basis of antibiotic resistance is complicated, and various mechanisms can be involved such as biofilm formation, pathway inhibition, degradation of antibiotics or regulation of efflux pumps^50–52^. In our study, we focused on serovar Typhimurium and discovered three iModulons related to cross-membrane transport that have increased activities for multidrug-resistant strains. It is also shown that chlortetracycline and florfenicol treatment represses the *crp* iModulon activity. The mechanism behind these patterns in response to antibiotic treatments can be further explored to assist drug and treatment designs in the future. Additionally, pan-genomic analysis identified flagella-related gene sets unique to the SPI-1 iModulon of the MDR strain D23580. With more data from MDR studies, it can be foreseen that our method has the potential to elucidate transcriptional patterns associated with drug resistance and guide experimental investigations.

In this study, ICA was demonstrated to be a great tool to understand the TRN of a bacterial species. The iModulons offered guidance in identifying MDR-related transport systems and new regulons related to *csgD* and tartrate utilization. iModulon activity clustering also elucidated CAMP resistance mechanisms and identified gene sets that act in concert to achieve specific biological functions. With more data, ICA has the potential to uncover the rich information still hidden in bacterial transcriptome data and encourage generation of more datasets to help reveal a more comprehensive TRN. Using pan-genomic analysis, we gained insights into similarities and differences in the TRN of related Typhimurium strains. We discovered genetic features related to differences in transcriptional and virulence signatures of the strains. This allowed for better understanding of the serovar as a whole and in specific strain, while linking genetic signatures to phenotypic differences in strains. As presented in this project, the combination of the two methods shows substantial promise towards enhancing our knowledge of the pathogen *Salmonella* to cope with this public health threat.

In addition to the results mentioned in this article, the complete results including ones not detailed here can be reproduced at https://github.com/AnnieYuan21/modulome-Salmonella. They are also available for the examination by researchers interested in *Salmonella* Typhimurium at an interactive portal https://imodulondb.org/.

## Methods

The complete code to reproduce this pipeline can be found at https://github.com/AnnieYuan21/modulome-Salmonella. The ICA part of the workflow is adapted from *Sastry*, A. V. *et al*. 2021^13^, which is also available at https://github.com/avsastry/modulome-workflow/.

### Pangenome Analysis and Phylogenetic Tree

We gathered the metadata of 3,329 *Salmonella* Typhimurium genomes from PATRIC^14^. To ensure genome quality, we removed genomes with less than 10x coverage and more than 150 contigs. Then 500 genomes were selected randomly from all the 1,199 Typhimurium genomes that passed quality control. We checked to include the strains that we had RNA-Seq data for which results in a final 506 strains. Using CD-HIT^15^ with a threshold of 0.9, we assembled the pan-genome matrix of these 506 strains. To identify the most “representative” strains, we used k-means clustering, and identified the optimal number of clusters with both explained variance and silhouette scores. With a silhouette score of 0.127 and explained variance of 86%, 177 clusters were chosen, giving 177 centroid strains. These 177 strains were then used to construct the phylogenetic tree (generated with snippy^54^ and gubbins^55^ with default parameters, using LT2 genome as a reference) and the Typhimurium core genome. Since some of the genomes of these strains were incomplete, we performed a sensitivity analysis to identify the size of the core genome. In the end, we used the cutoff at 4209 clusters from 172 genomes (**Figure 1b**). These clusters were defined in the PATRIC locus tags. To match with the locus tags of the expression profiles, PATRIC locus tags were mapped to the RefSeq LT2 locus tags. Some gene families were lost due to missing mapping between the two types of identifies, and the final core genome consists of 3,886 genes in RefSeq LT2 locus tag. The expressions of the core genome were extracted from the datasets of each strain and concatenated together to form the final core genome expression profile with 3,886 genes and 533 samples.

### RNA Seq Data Acquisition, Metadata Curation and Preprocessing

Following the PyModulon workflow (https://github.com/avsastry/modulome-workflow/tree/main/1_download_metadata), we compiled all of the RNA-seq data for *Salmonella* Typhimurium on NCBI SRA as of August 20, 2020. We performed manual curation of experimental metadata by inspecting literature associated with specific BioProject IDs documented in the metadata files. This is to identify different strains in the Typhimurium serovar and experimental conditions such as number of biological replicates, culture media, growth phase, temperature or any additional treatment that can assist us in understanding the results. Detailed metadata also helps with subsequent quality control steps. There were 1049 samples with detailed metadata, and in total 8 different Typhimurium strains in the datasets. Six strains had enough samples for subsequent analysis. Two strains (DT2 and sg_wt7) only had four samples each, so they were marked in the phylogenetic tree in **Figure 1a** and were used to construct the core genome, but were not investigated individually with ICA decomposition. The rest of the samples were then sorted into 6 separate files by strain, and processed using the RNA-Seq pipeline available at https://github.com/avsastry/modulome-workflow/tree/main/2_process_data. The final expression profiles were reported in units of log-transformed Transcripts per Million (log-TPM).

### Quality Control and Data Normalization

The expression datasets were then subject to five quality control steps outlined in https://github.com/avsastry/modulome-workflow/tree/main/3_quality_control. Briefly, we checked four statistics from the FastQC report, which are per base sequence quality, per sequence quality scores, per base n content, and adapter content and discarded all samples that didn’t pass these four criteria. Then, we removed samples with less than 5×10^5^ reads mapped to coding sequences. After that, we clustered the samples using hierarchical clustering and removed samples that did not conform to a typical expression profile, as these samples often use non-standard library preparation methods, such as ribosome sequencing and 3’ or 5’ end sequencing^56^. Then, samples with poor replicate correlation (Pearson R score < 0.9) and no biological replicates were discarded. However, there was an interesting project where biofilm formation was observed over a time course (PRJNA280002), so these samples were kept with a lower Pearson R score (0.8) to study temporal activities of iModulons. Also, PRJNA490148 contains samples from comparative studies of two Typhimurium strains. We were interested to see the differences in iModulon activities for two strains in the same conditions, so we included samples from this project even though some of them do not have biological replicates. After quality control, 533 samples were kept for ICA analysis.

To obviate any batch effects resulting from combining different expression profile datasets, we selected a reference condition in each project to normalize each dataset. This ensured that nearly all independent components generated were due to biological variation rather than technical variation. This normalization allows us to compare gene expression and iModulon activities within a project to a reference condition, but not across projects.

Six individual datasets were generated using the methods described above. To obtain the core genome expression profile, all gene locus tags were unified by using the LT2 locus tags from refseq, then the expression profiles were combined together based on the core genome that we defined earlier. The final core genome expression profile consists of 533 samples from 46 BioProjects distributed across 6 Typhimurium strains (**Figure 1c**).

### Defining the Optimal Number of Independent Components

To compute the optimal independent components, an extension of ICA was performed on the RNA-seq dataset as described in McConn et al.^57^.

Briefly, the scikit-learn (v0.23.2)^58^ implementation of FastICA^59^ was executed 100 times with random seeds and a convergence tolerance of 10^−5^. The resulting independent components (ICs) were clustered using DBSCAN^60^ to identify robust ICs, using an epsilon of 0.1 and minimum cluster seed size of 50. To account for identical with opposite signs, the following distance metric was used for computing the distance matrix:

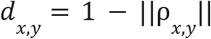

where *ρ*_*x,y*_ is the Pearson correlation between components *x* and *y*. The final robust ICs were defined as the centroids of the cluster.

Since the number of dimensions selected in ICA can alter the results, we applied the above procedure to each expression profile multiple times, ranging the number of dimensions from 5 to 380. Depending on the number of dimensionalities being tested, the step size varies from 5 (for smaller datasets with fewer dimensions) to 10 or 20 (for large datasets). The upper limit of the choice of dimensionality is approximately the number of samples in the dataset.

To identify the optimal dimensionality, we compared the number of ICs with single genes to the number of ICs that were correlated (Pearson R > 0.7) with the ICs in the largest dimension (called “final components”). We selected the number of dimensions where the number of non-single gene ICs was equal to the number of final components in that dimension (Supplementary Figure 8). The optimal number of dimensions for the core genome was 220. The optimal dimensionalities of the strains are presented in (Supplementary Figure 9).

ICA produces two matrices. The **M** matrix contains the robust independent components, and the **A** matrix contains the corresponding activities. The product of the **M** and **A** matrices approximates the expression matrix (the **X** matrix), which is the curated RNA-seq compendium. Each independent component in the **M** matrix is filtered to find the genes with the largest absolute weightings, which ultimately generates gene sets that make up iModulons. Implementing this process on the Typhimurium core genome resulted in 115 iModulons that explained 75% of the expression variance in the core compendium (Supplementary Figure 2). The number of iModulon from the strain expression profiles can be found in **Figure 5a.**

### Compiling TRN and Gene Annotations

The TRN file was generated using a pre-defined TRN for the closely related bacteria *Escherichia coli*^18^ and the bi-directional blast results of Typhimurium LT2 and *E. coli* genomes https://github.com/AnnieYuan21/modulome-Salmonella. The genes are annotated following the annotation pipeline that can be found at https://github.com/SBRG/pymodulon/blob/master/docs/tutorials/creating_the_gene_table.ipynb. Additionally, KEGG^61^ and Cluster of Orthologous Groups (COG) information were obtained using EggNOG mapper^62^. Uniprot IDs were obtained using the Uniprot ID mapper^63^, and operon information was obtained from Biocyc^64^. Gene ontology (GO) annotations were obtained from AmiGO2^65^.

### Computing iModulon Enrichments

iModulon enrichments against known regulons were computed using two-sided Fisher’s exact test, with the false discovery rate (FDR) controlled at 10^−5^ using the Benjamini-Hochberg correction. Functional enrichment through KEGG and GO annotations were similarly computed but with FDR < 0.01. This pipeline can be found at https://github.com/SBRG/pymodulon/blob/master/docs/tutorials/gene_enrichment_analysis.ipynb.

### Differential iModulon Activity Analysis

The differences in iModulon activities under relevant conditions were calculated using a log normal probability distribution. For each comparison, the absolute difference in the mean iModulon activities were calculated and compared to an iModulon’s log-normal distribution (calculated between biological replicates). P-values statistics were obtained for each condition comparison across all iModulons and a FDR score was calculated. iModulons with a difference greater than 5 and FDR less than 0.01 are considered significant. Differential iModulon activity plot (DIMA plots) can be generated to visualize iModulon activity differences under one condition against another.

### Calculating iModulon Activity Clusters

The activities of iModulons were clustered using the Seaborn^66^ *clustermap* function in Python. The default distance metric is the following:

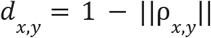

where ||ρ_*x,y*_|| is the absolute value of the Spearman R correlation between two iModulon activity profiles. Other available distance metric options include Pearson R, Kendall Rank and Mutual Information. The option used for this project was Mutual Information. The threshold for optimal clustering was determined by testing different distance thresholds to locate the maximum silhouette score.

### Generating iModulonDB Dashboards

iModulonDB dashboards were generated using the PyModulon package^13,67^; the pipeline can also be found at https://pymodulon.readthedocs.io/en/latest/tutorials/creating_an_imodulondb_dashboard.html.

## Supporting information

Supplementary File 1_Core expression profile

Supplementary File 2_Core metadata

