## Supplementary Notes for "Pan-genomic analysis of transcriptional modules across *Salmonella* Typhimurium reveals the regulatory landscape of different strains"

### Supplementary Note 1

It was documented that Typhimurium strain ATCC 14028 was once termed just as strain 14028. However, a derivative of this original strain was found to have rough colony morphology, so it was named as 14028r (r for rough)[1]. To avoid confusion, the original smooth strain was then renamed as 14028s. This record seems to suggest that these three strains are in fact the same. Looking at the phylogenetic tree, the three strains are found in the same node close to each other, confirming their relatedness. Therefore, in our downstream analysis, these three strains were combined together and named as 14028S.

### Supplementary Note 2

The name used in the original bioproject is "tartarate" and we assumed it meant tartrate[2]. We are not sure which tartrate isomer was used (L/D/meso) or the aerobicity of the experiments.
