## Supplementary figures and images for "Pan-genomic analysis of transcriptional modules across *Salmonella* Typhimurium reveals the regulatory landscape of different strains"

### Supplementary Figure 1

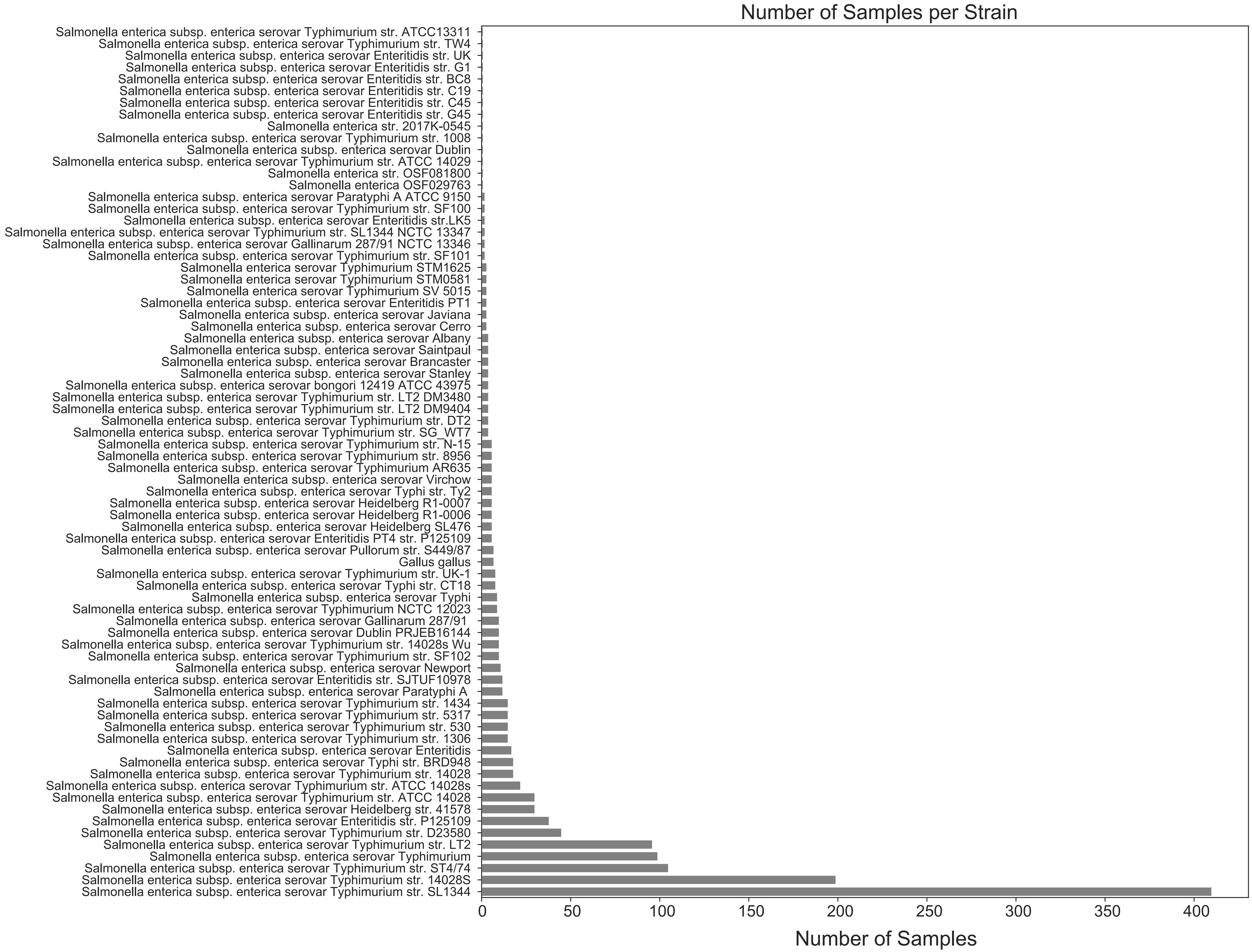

### Supplementary Figure 2

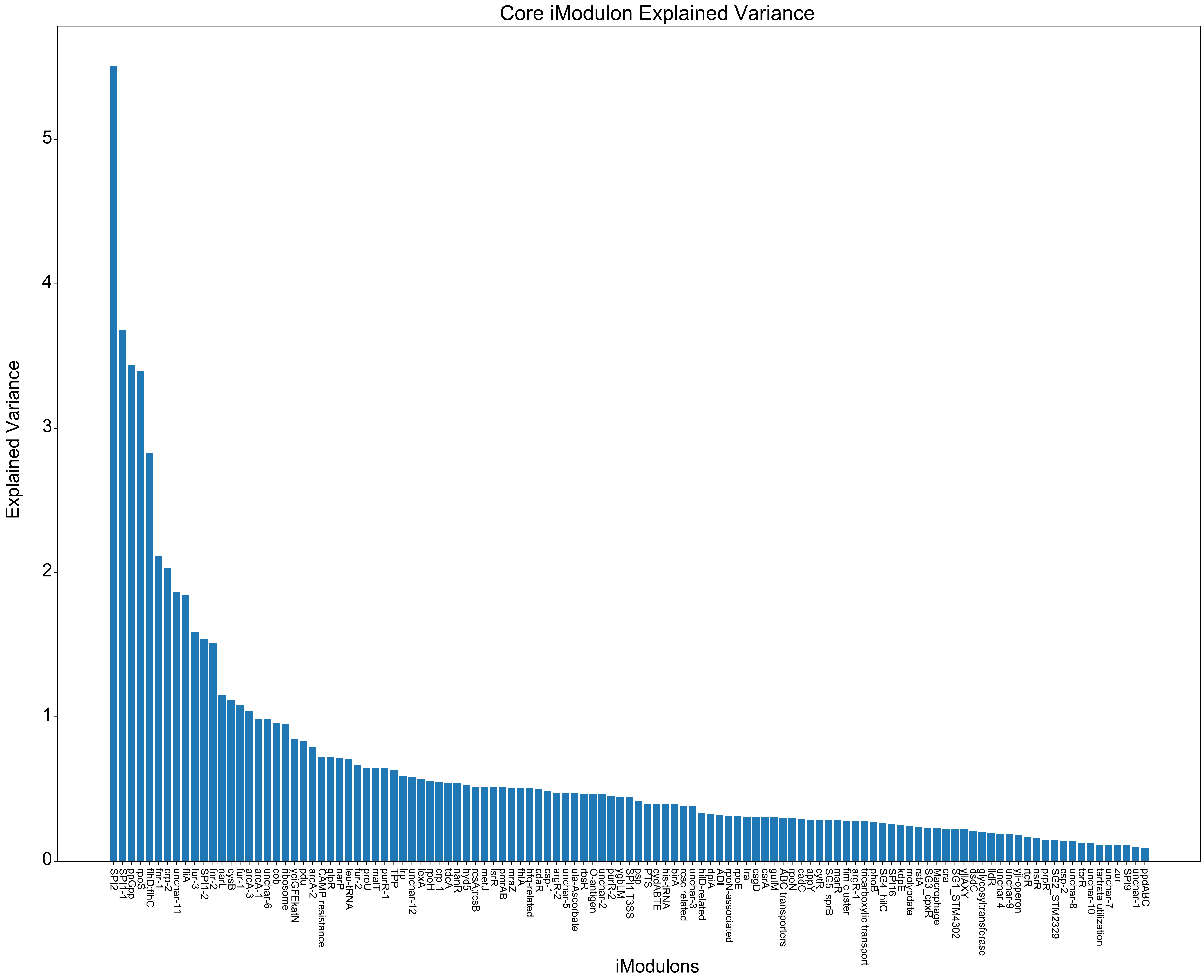

### Supplementary Figure 3

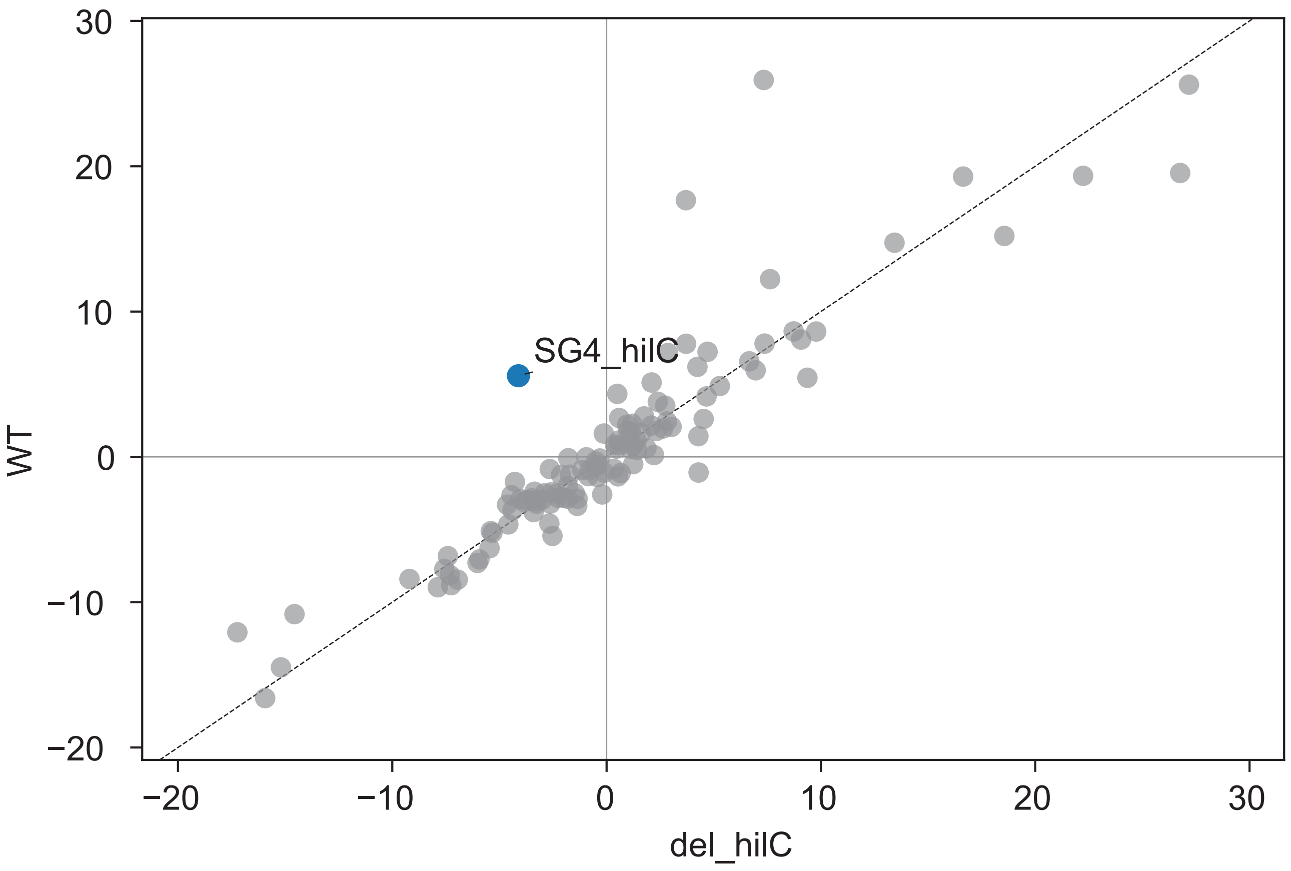

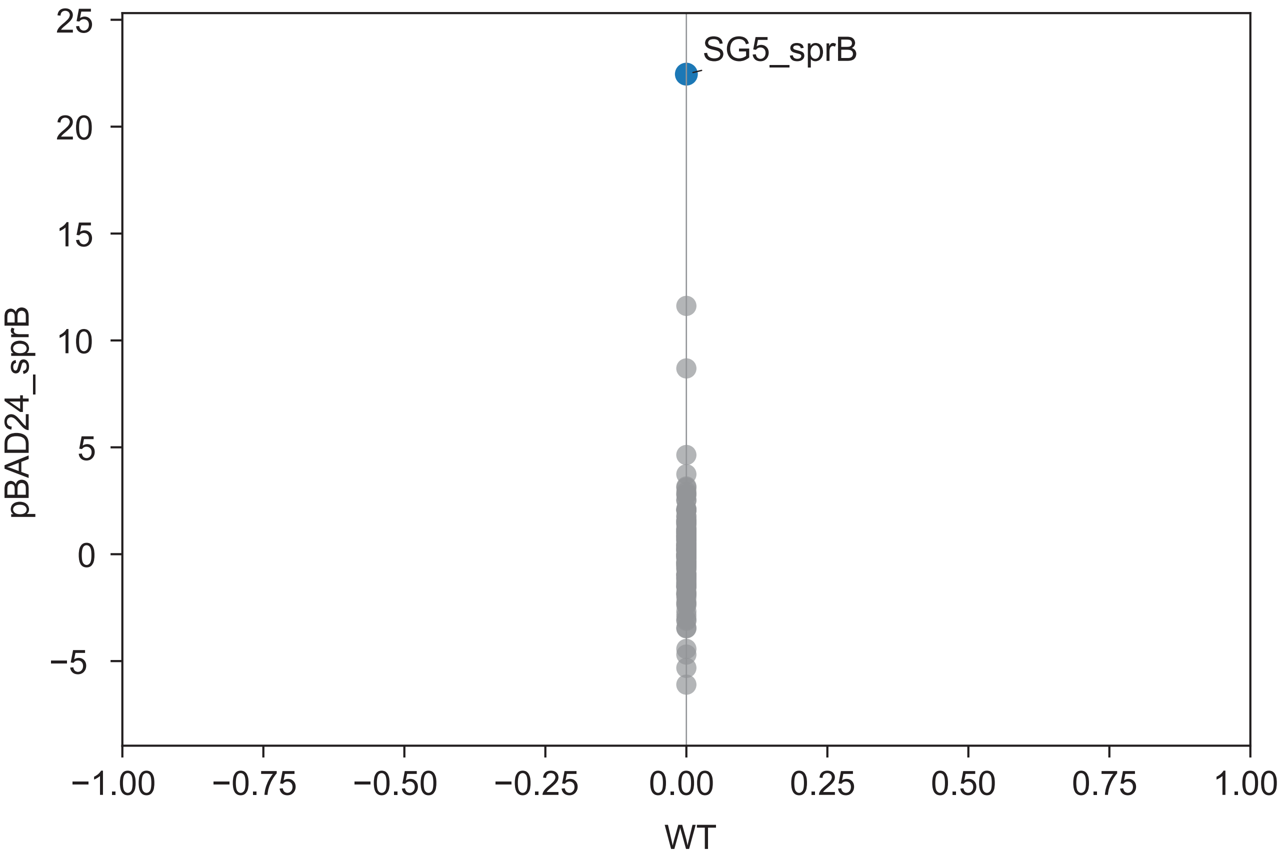

### Supplementary Figure 4

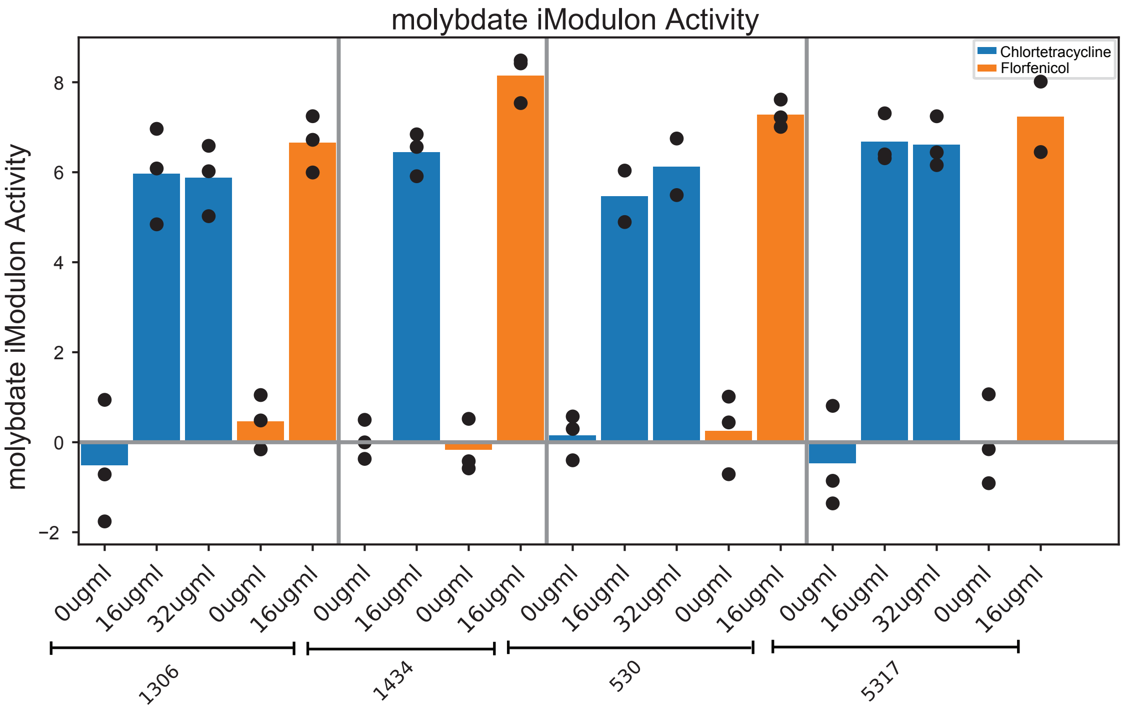

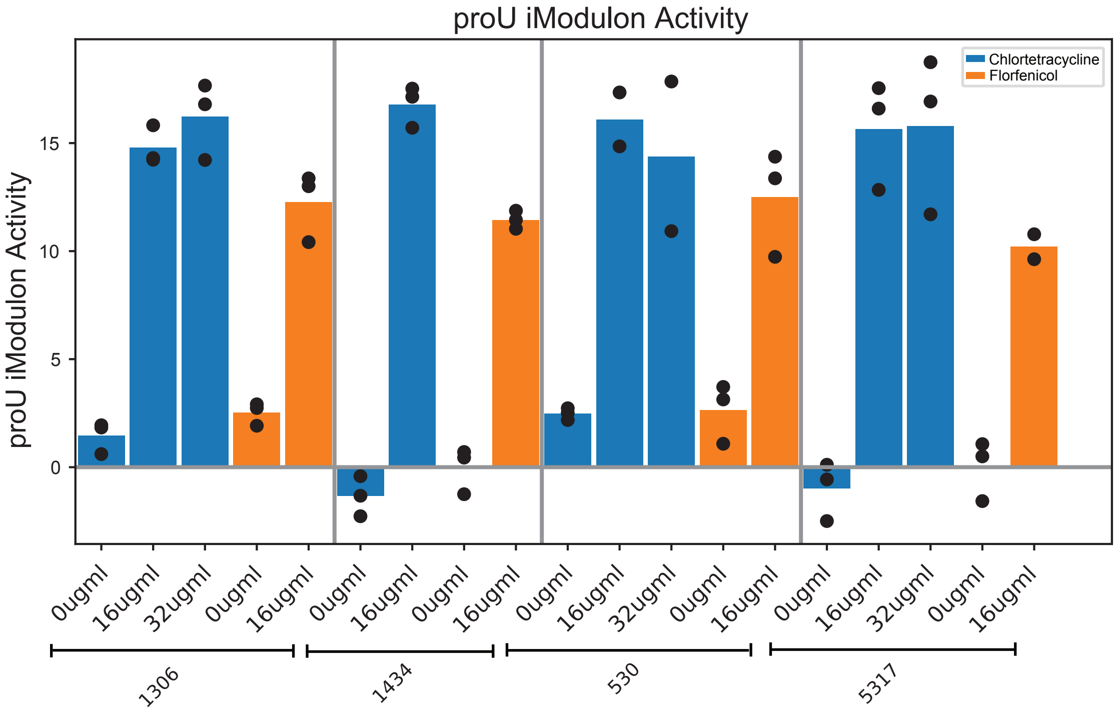

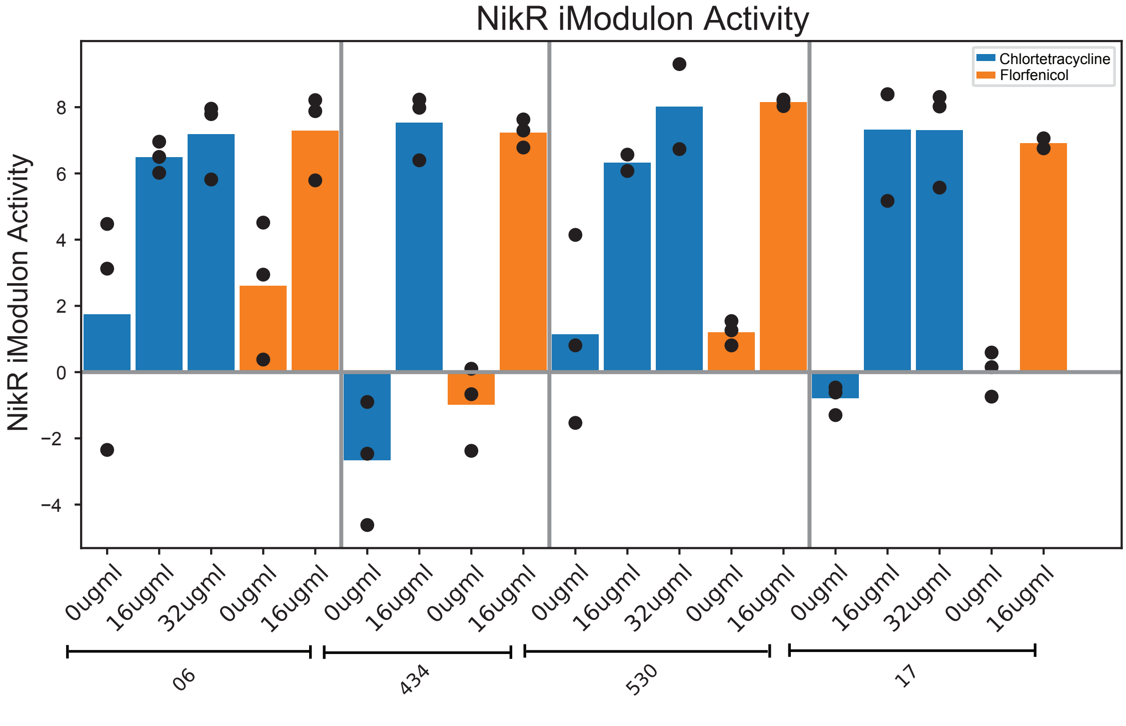

### Supplementary Figure 5

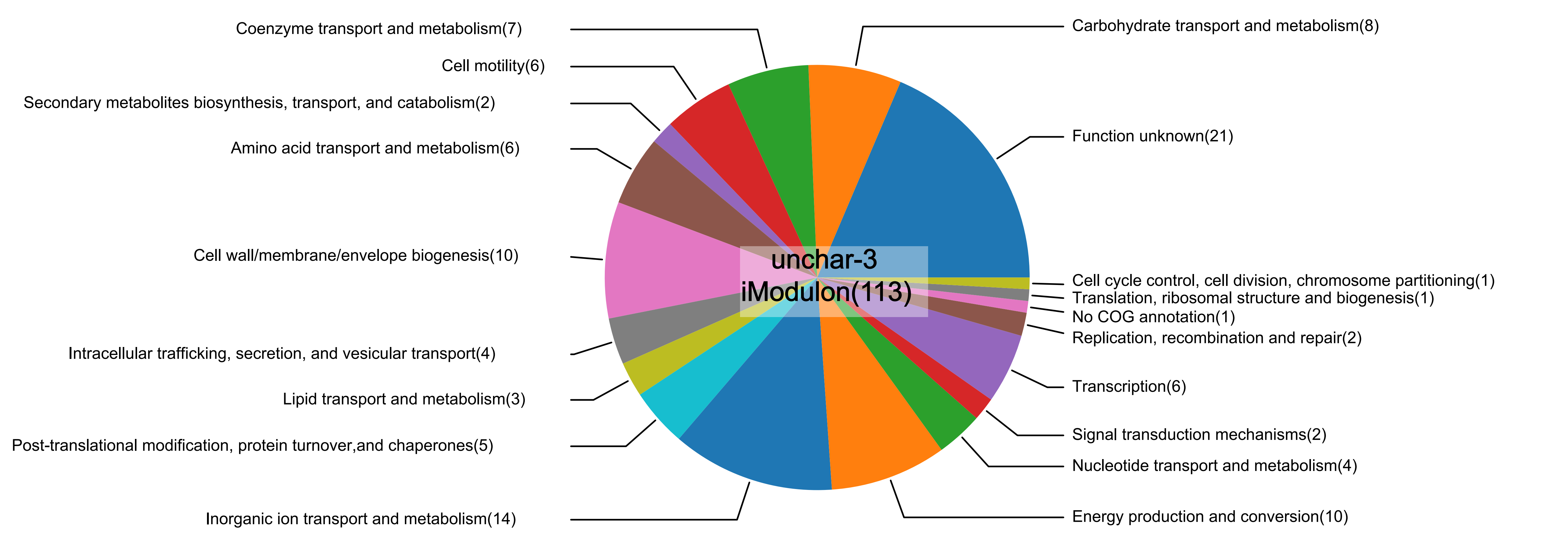

### Supplementary Figure 6

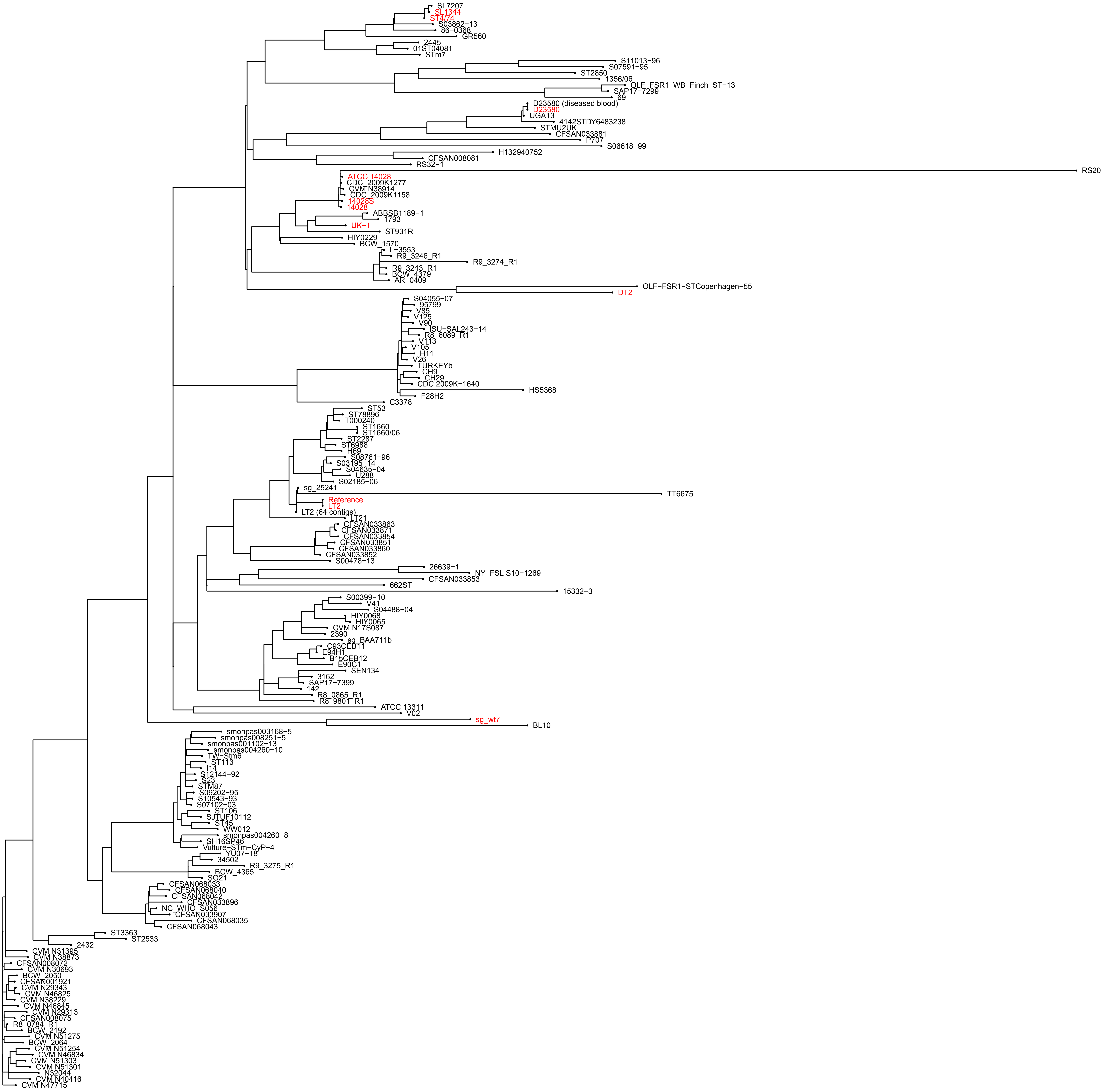

### Supplementary Figure 7

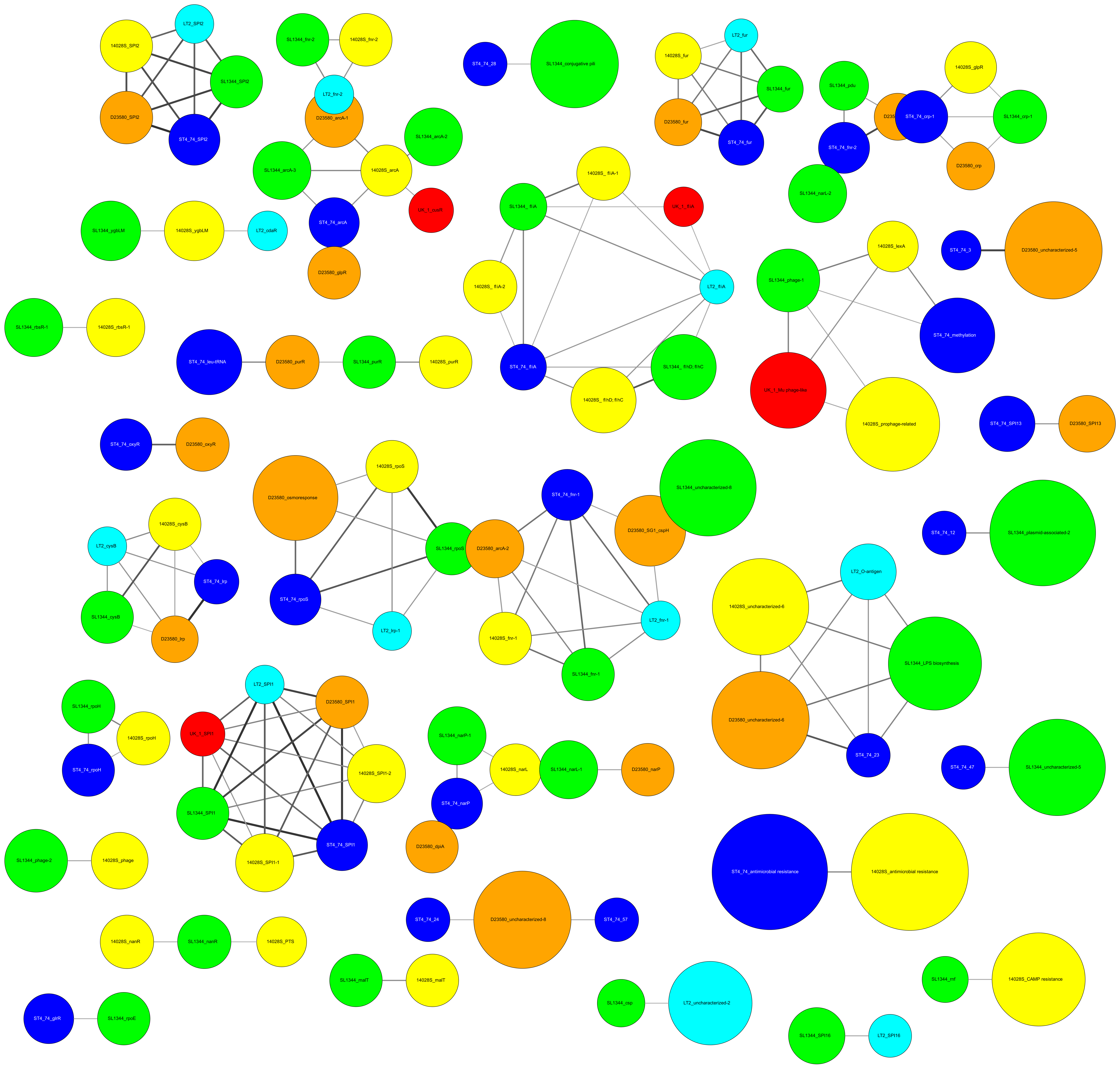

### Supplementary Figure 8

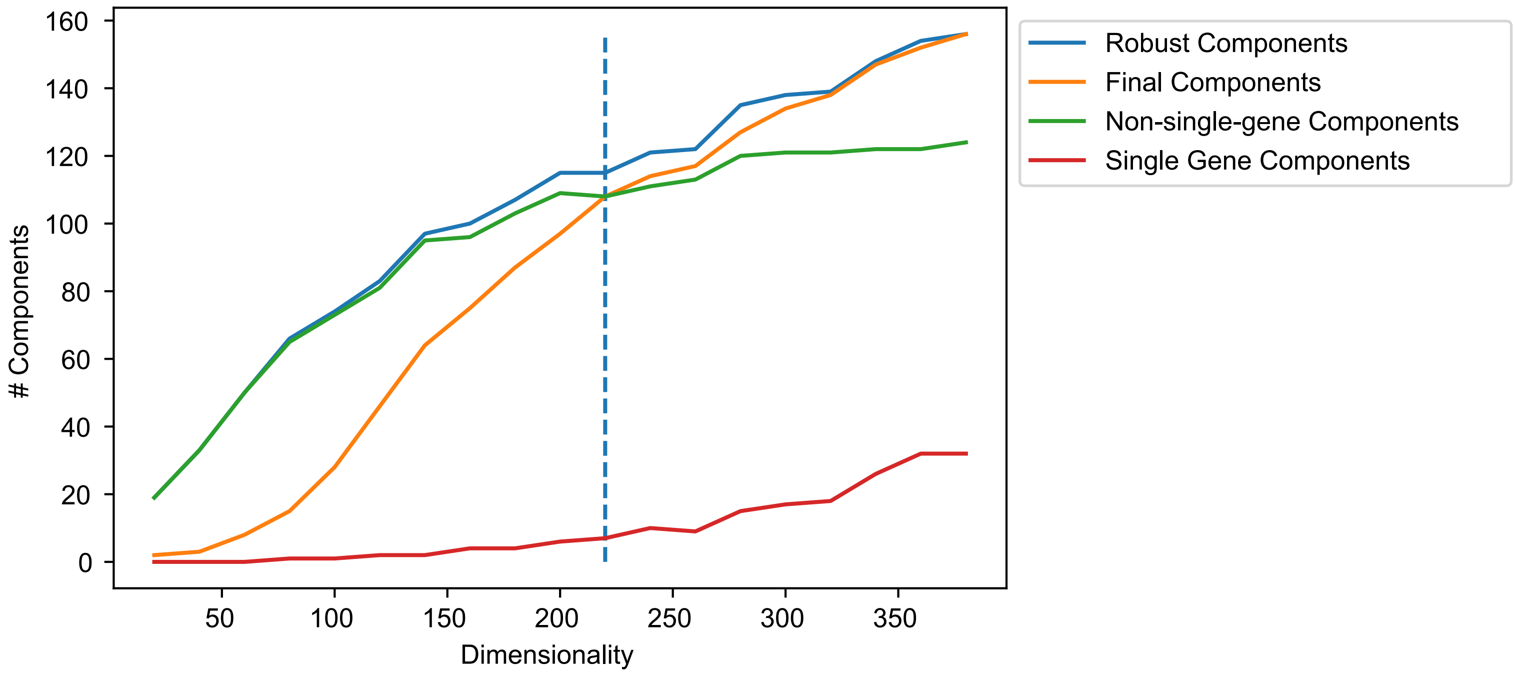

### Supplementary Figure 9

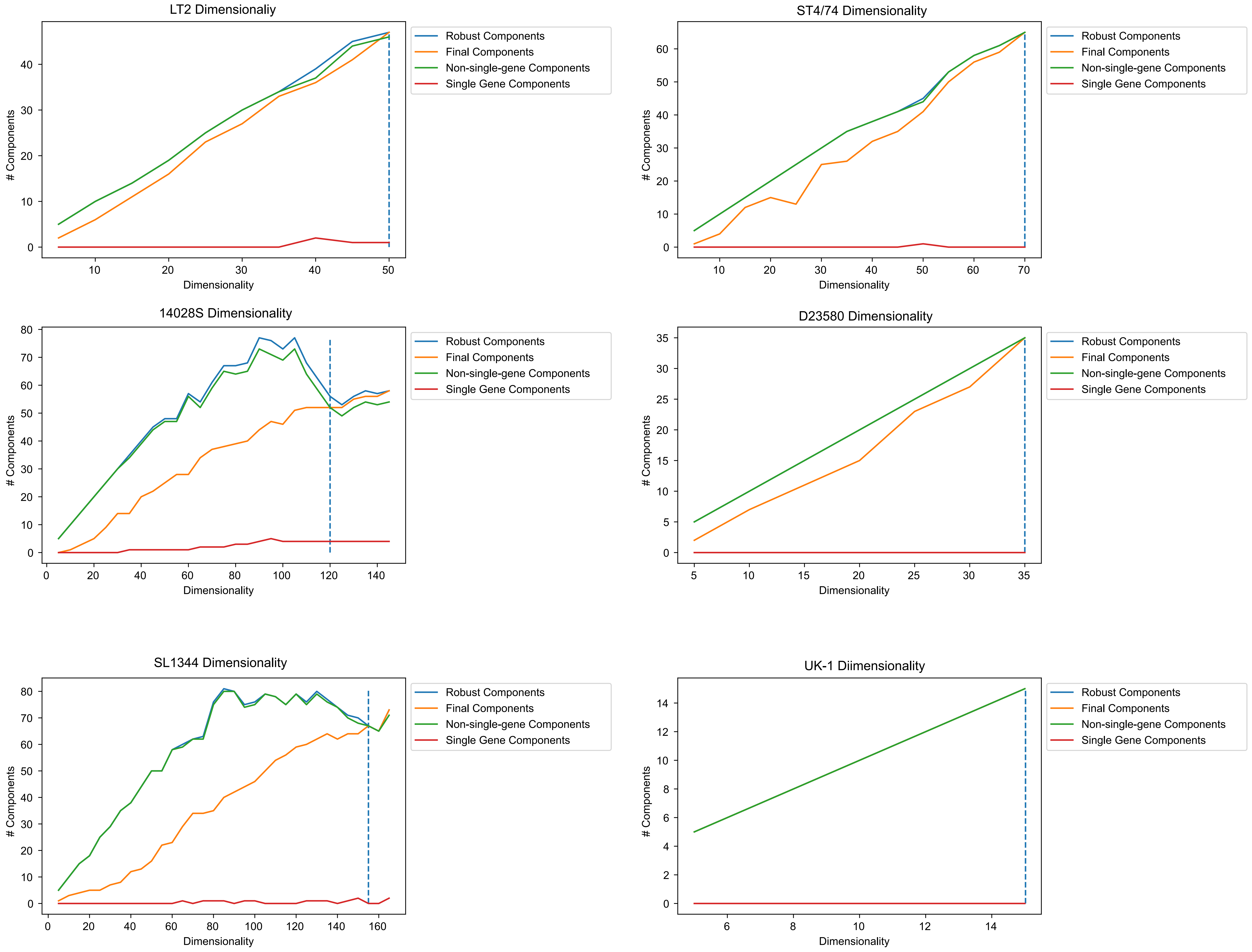
